# A helper NLR channels organellar calcium to trigger plant immunity

**DOI:** 10.1101/2024.09.19.613839

**Authors:** Tarhan Ibrahim, Freddie J. King, AmirAli Toghani, Luyao Wang, Saskia Jenkins, Enoch Lok Him Yuen, Hung-Yu Wang, Cristina Vuolo, Nick Eilmann, Vanda Adamkova, Khong-Sam Chia, Baptiste Castel, Jonathan D. G. Jones, Philip Carella, Chih-Hang Wu, Jiorgos Kourelis, Sophien Kamoun, Tolga O. Bozkurt

**Affiliations:** Department of Life Sciences, Imperial College London; London SW7 2AZ, UK; The Sainsbury Laboratory, University of East Anglia; Norwich Research Park, Norwich NR4 7UH, UK; Institute of Plant and Microbial Biology, Academia Sinica; Taipei 115201 Taiwan; Cell and Developmental Biology, John Innes Centre; Norwich NR4 7UH, UK

**Author notes:** Corresponding authors (T.O.B.), (J.K.) and (S.K.). These authors contributed equally to this work (Tarhan Ibrahim, Freddie J. King).

## Abstract

Upon activation, plant NLR immune receptors are known to assemble into oligomeric resistosomes that insert into the plasma membrane, forming Ca^2+^-permeable channels and triggering immunity. Here, we show that the RPW8-like coiled-coil (CC_R_-) NLR NRG1 primarily targets organelles instead of the plasma membrane. Unlike canonical CC-NLRs, activated NRG1 accumulates at the chloroplast envelope and channels stromal Ca^2+^ into the cytosol. AlphaFold modeling of the NRG1 resistosome reveals an unusually long N-terminal membrane-insertion structure that would span the double chloroplast membranes. Nanobody-mediated relocalization shows functional membrane specificity: chloroplast trapping abolishes activity of the canonical helper CC-NLR NRC4 but not NRG1. NRG1 orthologs, from non-flowering lineages to angiosperms, target chloroplasts, suggesting that organelle-centered defense dates back to at least ∼360 million years. We propose that coiled-coil NLR diversification has enabled compartment-specific immune signaling to capture diverse Ca^2+^ stores.

## Main Text

Innate immunity relies on the precise recognition of pathogens by nucleotide-binding leucine-rich repeat receptors (NLRs), which coordinate defense-related programmed cell death (*1–2*). In mammals, activated NLRs assemble into inflammasomes that direct Gasdermin oligomers to the plasma membrane, leading to pyroptotic cell death and pathogen clearance (*3–4*). Similarly, in plants, activated NLRs assemble into oligomeric structures called resistosomes, some of which target the plasma membrane, ultimately leading to a hypersensitive response (HR) cell death, thereby restricting pathogen invasion (*5–8*).

Some NLRs function as singleton receptors to both sense and signal the presence of pathogen effectors. Others operate in specialized pairs or complex networks, where sensor NLRs recognize pathogen effectors and helper NLRs execute the HR. NLRs are multidomain receptors that typically consist of a divergent N-terminal domain, a central NB-ARC domain, and a C-terminal leucine-rich repeat (LRR) domain. Phylogenetic classification divides flowering plant NLRs into four major classes, each characterized by specific N-terminal domain architectures: Toll/Interleukin-1 Receptor (TIR)-NLRs and three types of sequence-unrelated but structurally and functionally conserved 4-helical bundle coiled-coil domains: canonical coiled-coil (CC) domains, some of which carry a conserved MADA motif (e.g., ZAR1 and NRCs), RPW8-like domains characteristic of CC_R_-NLRs (e.g., NRG1 and ADR1), or G10-type domains (CC_G10_-NLRs; e.g., RPS2 and RPS5). In many canonical CC-NLRs, activation triggers oligomerization into resistosomes that function as Ca^2+^-permeable pores, initiating immunity (*9–12*).

Canonical CC-NLRs (ZAR1, Sr35) oligomerize into pentameric resistosomes, while CC helper NLRs from the NRC clade (NRC2/4) form hexameric resistosomes, both targeting the plasma membrane (*8, 13–18*). By contrast, TIR-NLRs assemble into tetrameric resistosomes that produce chemical signals indirectly activating downstream CC_R_-NLR helpers like NRG1 and ADR1 (*19–21*). Although CC_R_-NLRs form oligomers with calcium channel activity similar to canonical CC-NLR resistosomes, complete structures of CC_R_-NLR resistosomes remain unresolved despite recent partial structures of monomeric NRG1 (*6, 22–25*). However, another common feature of the activated CC-NLRs and CC_R_-NLRs is localization to distinctive punctate structures in the cell, likely representing clusters of resistosomes, which can be observed through confocal microscopy using C-terminally fused fluorescent tags (*15, 18, 26*).

The N-terminal four-helix bundles that execute cell death in plant NLRs are structurally analogous to pore-forming domains in other kingdoms, including mammalian Mixed Lineage Kinase domain-Like (MLKL) proteins and fungal heterokaryon incompatibility proteins (*6, 27–29*). Despite this conserved architecture, the amino acid sequences of NLR N-terminal domains that execute immunity are remarkably diverse in plants (*17, 28*). This paradox suggests that resistosome functions may extend beyond the established paradigm of acting as plasma-membrane Ca^2+^ channels. We therefore investigated the CC_R_-NLR NRG1, whose divergent N-terminus provided an ideal system to investigate whether this architectural variation enables alternative modes of immune activation.

### NbNRG1 localizes to multiple subcellular membranes

To understand the activation mechanism of NRG1, we first examined the sub-cellular localisation of NRG1 from the solanaceous model plant *Nicotiana benthamiana* (NbNRG1) in both resting and activated states. Transient expression of full-length NbNRG1:GFP in *N. benthamian*a causes cell death (Fig. S1) (*30*), hampering live-cell imaging and other cell-biology analyses. To reduce the auto-activity of NbNRG1, we generated a truncated clone of NbNRG1 lacking the first fourteen amino acids of the protein, the most N-terminal region of the CC_R_ domain (NbNRG1^Δ14^). This deletion is equivalent to that previously used for studying *Arabidopsis thaliana* NRG1.1 (AtNRG1.1) (Data S1) (*6, 25*). Transiently expressing NbNRG1^Δ14^:GFP triggered weaker cell death compared to full-length NbNRG1:GFP, making it more suitable for cell biology study (Fig. S1). Importantly, NbNRG1^Δ14^ retained the ability to be activated and form punctate structures. Activation occurred upon co-expression with the silencing suppressor p19, which increases transient expression levels, or when activated by the *Xanthomonas* effector XopQ, which is recognised by the TIR-NLR Roq1 in *N. benthamiana* (*31, 32*). While monitoring the subcellular localisation of NbNRG1^Δ14^:GFP, we observed atypical punctate fluorescence rather than the uniform peripheral punctate pattern characteristic of plasma-membrane localisation, suggesting that activated NbNRG1 may target other membranous compartments. To test for organelle targeting, we co-expressed NbNRG1^Δ14^:GFP with RFP-tagged organelle markers. NbNRG1^Δ14^:GFP puncta co-localised with markers for the endoplasmic reticulum (ER) and mitochondria, but not with Golgi or plasma-membrane markers, nor with a free-RFP control (Fig. 1, A to E). To exclude the possibility that the observed localization was an artifact of the truncated NbNRG1 variant, we optimized full-length NbNRG1 expression to mitigate cell death and imaged cells 26-28 hours after agroinfiltration. Full-length NbNRG1:mGOLD, like NbNRG1^Δ14^:GFP, localized to organellar compartments rather than the plasma membrane (Fig. S2). We next examined whether NbNRG1 activation and localization depends on the canonical XopQ-triggered EDS1-SAG101 signaling cascade. At 24 hours post infiltration, NRG1:mTurquoise2 expressed in *N. benthamiana nrg1/adr1* and *epss* knockout plants remained largely cytoplasmic when co-expressed with a GUS control. Under the same conditions, co-expression with XopQ induced prominent puncta around chloroplasts in *nrg1/adr1* plants, but not in *epss* plants. By 2 days post infiltration, XopQ co-expression triggered HR symptoms in *nrg1/adr1* but not *epss* plants. Together, these results suggest that the canonical EDS1-SAG101 signalling module is required for effector-dependent NRG1 activation and subsequent localization to chloroplast (Fig. S3). To determine whether organelle localisation is unique to the CC_R_-NLR NbNRG1, we monitored the subcellular localisation of an auto-active CC-NLR helper NbNRC4^D478V/L9E^:GFP using the same set of markers. Consistent with previous reports, NbNRC4^D478V/L9E^ puncta accumulated exclusively at the plasma membrane, with no punctate signal at subcellular membranes or the cytoplasm (Fig. S4) (*26*). Collectively, these results reveal that the helper CC_R_-NLR NbNRG1 undergoes a non-canonical trafficking route upon activation, localising to multiple endomembranes rather than the plasma membrane, in stark contrast to the canonical CC-NLR resistosomes (*8, 18, 26*).

**Fig. 1.**
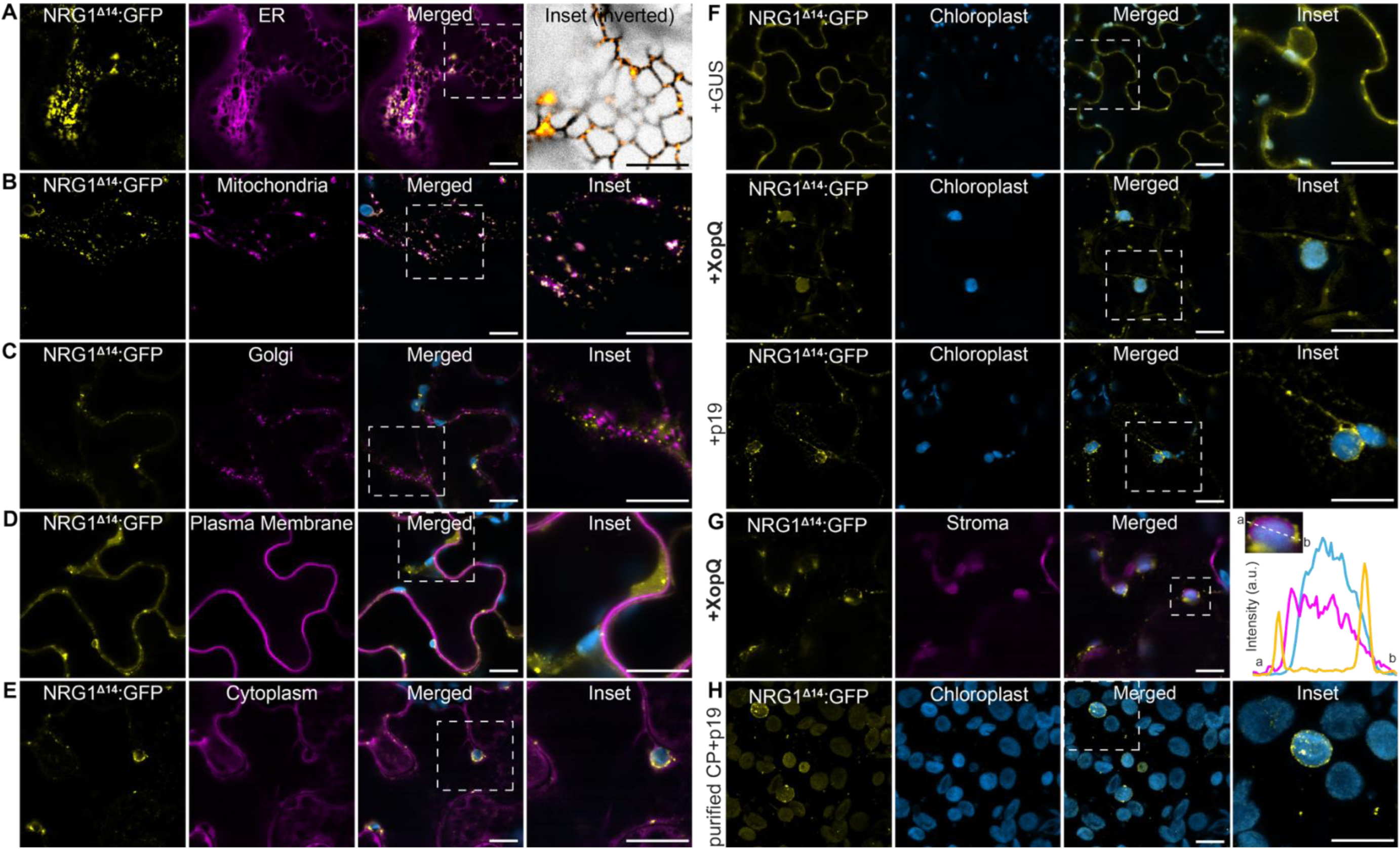
NbNRG1 targets various subcellular membranous compartments. Confocal micrograph of *N. benthamiana* leaf epidermal cells transiently expressing NbNRG1^Δ14^:GFP with (**A**) SP:RFP:HDEL (ER), (**B**) ScCOX41-29:mCherry (mitochondria), (**C**) GmMan11-49:mCherry (Golgi), (**D**) RFP:Remorin1.3 (plasma membrane) and (**E**) RFP:EV (Cytoplasm). Dashed lines in merged panels correspond to insets. (**F**) Confocal micrographs of *N. benthamiana* leaf epidermal cells transiently expressing NbNRG1^Δ14^:GFP with GUS, XopQ and p19. Dashed lines in merged panels correspond to insets. (**G**) Confocal micrograph of *N. benthamiana* leaf epidermal cells transiently expressing NbNRG1^Δ14^:GFP with XopQ and CTP1-RFP (Stroma). The merged panel highlights chloroplast autofluorescence. Dashed lines in the merged panel correspond to the line intensity plot depicting the relative fluorescence across the marked area. The lines in the intensity plot are colored accordingly with the fluorescent tags used. (**H**) Confocal micrograph of purified chloroplasts (CP) from *N. benthamiana* leaf tissue transiently expressing NbNRG1^Δ14^:GFP with p19. Dashed lines in the merged panel correspond to inset. Chloroplasts are visualized using auto-fluorescence at 648-709 nm. All images shown are single-plane images. Scale bars represent 10 μm. Imaging was performed at 2-days post-agroinfiltration using a minimum of two-leaf patches per sample.

### Activated NbNRG1 localizes to the chloroplast envelope

While examining the various organelle targets of NbNRG1, we noticed that a subset of puncta appeared adjacent to plastids. This prompted us to examine in more detail whether activated NbNRG1 associates with chloroplast membranes. We focused on chloroplasts because they are among the most easily discerned organelles in plant cells. In addition, given the availability of reliable markers, intrinsic autofluorescence that facilitates live-cell imaging, and established tools for functional studies, we focused subsequent analyses on the chloroplast as a tractable model organelle. To monitor NbNRG1 localisation in an active state, we co-expressed NbNRG1^Δ14^:GFP with XopQ or p19 and observed puncta localization with confocal microscopy. Active NbNRG1^Δ14^:GFP displayed a prominent punctate distribution with a subset of the puncta localizing to chloroplasts (Fig. 1F). In contrast, co-expression with the negative control GUS revealed that resting NbNRG1^Δ14^ is cytoplasmic (33% of cells, *n*=12), consistent with other helper NLRs (Fig. 1F and Fig. S5A) (*18, 26*). Even in the absence of XopQ, NbNRG1^Δ14^ retained partial auto-activity with chloroplast-associated puncta observed in 67% of cells (*n*=12). Upon further activation by XopQ, every examined cell displayed punctate localisation (11/11), and many puncta were chloroplast-associated (Fig. S5A). Using Blue-native PAGE, we confirmed that co-expression with XopQ shifts NbNRG1 into a higher-molecular-weight complex, consistent with activation and resistosome formation, linking functional resistosomes to puncta observed (Fig. S5D). To pinpoint the chloroplast localization of NbNRG1, we labelled the stroma with CTP1:RFP (*33*). NbNRG1^Δ14^:GFP puncta encircled the stromal marker, indicating that NbNRG1 engages with the chloroplast envelope (Fig. 1G). To confirm chloroplast envelope localization, we used TOC64:GFP as a marker for the outer chloroplast membrane (*34*). Full-length NbNRG1:mScarlet3 puncta co-localized with TOC64:GFP, demonstrating that NbNRG1 associates with the outer membrane (Fig. S6). We also detected NbNRG1:mScarlet3 puncta on stromules, the tubular extensions of chloroplast stroma (*35*). We further validated these live-cell imaging observations with purified chloroplasts, confirming the punctate distribution of NbNRG1^Δ14^ on the chloroplast surface (Fig. 1H).

To investigate the subcellular dynamics of resting state NbNRG1, we tracked its localization pattern under the weaker *Arabidopsis* Act2 promoter (pAct2) and with monomeric fluorescent tags. Unlike p35S::NbNRG1^Δ14^:GFP, pAct2::NbNRG1^Δ14^:GFP did not cause any visible cell-death symptoms (Fig. S5C). Consistent with the loss of cell death, pAct2::NbNRG1^Δ14^:GFP showed a weak cytoplasmic signal (78%, *n*=18 cells), with only few cells displaying detectable puncta, some of which were associated with chloroplasts (*n*=4/18 cells, Fig. S5B). To corroborate these results with full-length NbNRG1, we tagged it with the monomeric fluorescent protein mKOk (p35S::NbNRG1:mKOk) (*36*) which also caused weaker cell death compared to NbNRG1:GFP, enabling live cell imaging (Fig. S5C). NbNRG1:mKOk showed a mixed localization pattern in the few surviving cells. The protein displayed either cytoplasmic distribution (resting state) or punctate structures, some of which localized to chloroplasts in *nrg1* knockout plants (*n*=5/8 images, Fig. S7A). This pattern remained similar in *N. benthamiana nrg1/adr1* double knockouts upon co-expression with XopQ or a GUS negative control (Fig. S7B). Notably, NbNRG1:mKOk remained capable of activation by co-expression with XopQ, forming puncta more frequently (75% of cells, *n*=16/20 images) compared to the negative control (30% of cells, *n*=6/20 images). Similarly, activation of full-length NbNRG1 expressed from the weaker pAct2 promoter and fused to the bright td8ox2StayGold fluorophore (*37*) produced a prominent punctate pattern, with a subset of puncta surrounding the chloroplasts (Fig. S7C). Overall, using full-length and truncated NbNRG1 with multiple fluorophores and various promoters, in several genetic backgrounds, we have shown that resting state NbNRG1 is largely cytoplasmic but, upon activation, NbNRG1 can localize to the chloroplast envelope.

### NbNRG1 facilitates calcium efflux from chloroplasts

AtNRG1.1 can oligomerize, form puncta, and channel Ca^2+^, common features of helper NLRs (*6, 16*). We first established that NbNRG1 elevates cytosolic Ca^2+^, consistent with the prior reports (*6*) (Fig. S8). To determine whether NbNRG1 fulfils a similar role at chloroplasts, we co-expressed full-length NbNRG1 fused to mScarlet3 (NbNRG1:mScarlet3) with a stromal Ca^2+^ sensor. We used the genetically encoded calcium sensor GCaMP6s, targeted to the stroma with the RBCS1A transit peptide (RBCS1A-GCaMP6s) (*38, 39*). Chloroplasts with NbNRG1 puncta displayed markedly lower GCaMP6s fluorescence than neighbouring chloroplasts lacking puncta, indicating reduced stromal Ca^2+^ levels (Fig. S9). To monitor calcium flux from the chloroplast in the absence of cell death, we deleted the first 20 amino acids of the CC_R_ domain of NbNRG1 (NbNRG1^Δ20^). The NbNRG1^Δ20^ truncation completely abolished cell-death symptoms compared to full-length NbNRG1 and NbNRG1^Δ14^, yet preserved chloroplast localisation (Fig. 2A and Fig. S10). To determine if NbNRG1 requires the N-terminus of the CC domain to channel Ca^2+^ from the chloroplast, we co-expressed the stromal Ca^2+^ sensor (RBCS1A-GCaMP6s) with either NbNRG1:mScarlet3, NbNRG1^Δ20^:mScarlet3, or GUS:mScarlet3, and quantified the relative sensor fluorescence. Our quantitative fluorescence measurements revealed that chloroplasts with full-length NbNRG1:mScarlet3 puncta displayed the greatest reduction in stromal GCaMP6s signal, significantly lower than that seen with either the NbNRG1^Δ20^ or the GUS:mScarlet3 control (Fig. 2, A and B). To examine Ca^2+^ flux upon XopQ activation, we co-expressed full-length NRG1:mTurquoise2 with chloroplast (RBCS1A-GCaMP6s) and cytoplasmic (RCaMP1h) Ca^2+^ sensors (*40*). Activation by XopQ induced NRG1:mTurquoise2 puncta on the chloroplast envelope and a marked increase in the RCaMP1h:GCaMP6s fluorescence ratio, consistent with Ca^2+^ efflux from the chloroplast into the cytoplasm relative to the GUS control (Fig. 2, C and D).

**Fig. 2.**
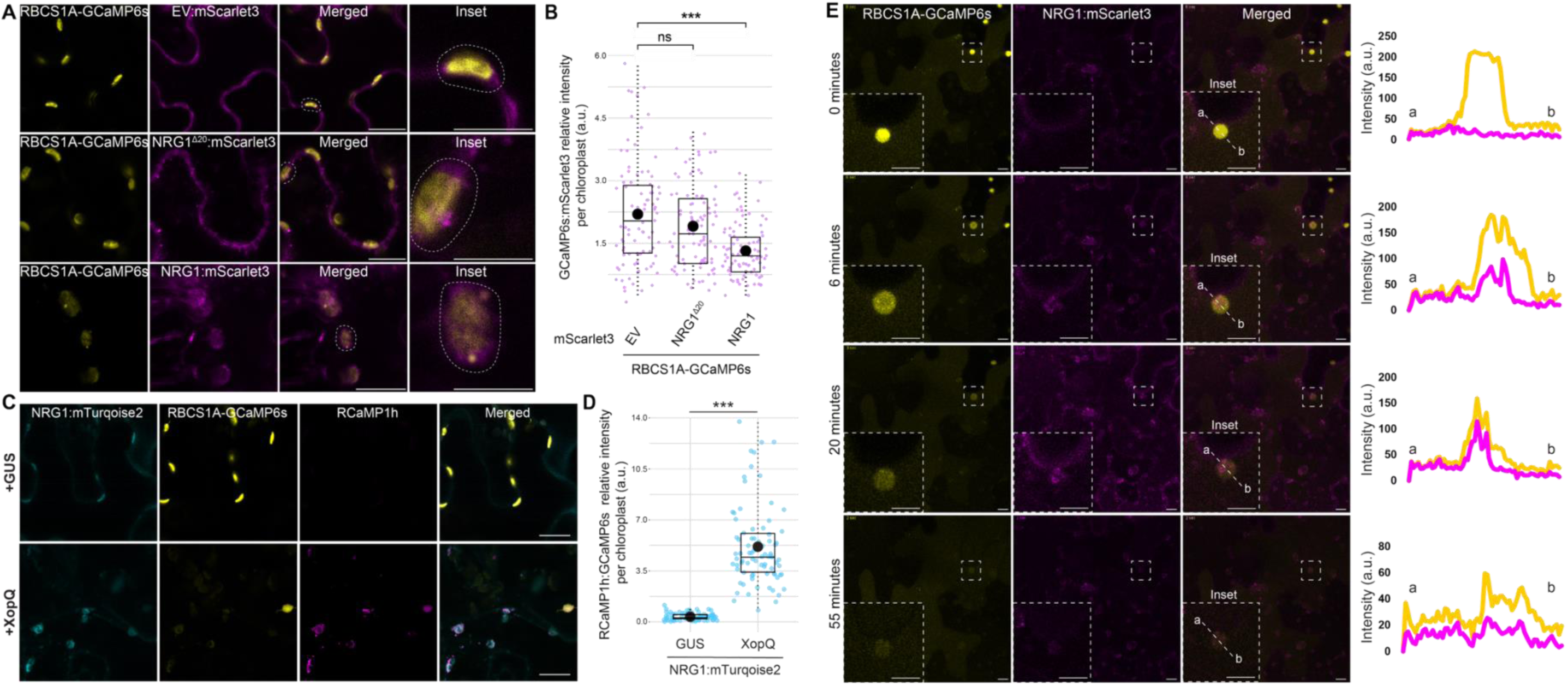
NbNRG1 facilitates calcium ion efflux from chloroplasts. (**A**) Confocal micrographs of *N. benthamiana* leaf epidermal cells transiently expressing RBCS1A-GCaMP6s:GFP with EV:mScarlet3, NRG1^Δ20^:mScarlet3 or NRG1:mScarlet3. Dashed lines in merged panels correspond to inset, which demonstrates examples of chloroplasts used to measure relative GCaMP6s:mScarlet3 intensity in (B). All images shown are single-plane images. Scale bars represent 10 μm. (**B**) Box and dot plot showing the GCaMP6s:mScarlet3 relative intensity per chloroplast for 90 chloroplasts. Each chloroplast is highlighted with a lilac dot. The mean value for each condition is shown as a black dot. Asterisks indicate statistically significant difference of NRG1^Δ20^:mScarlet3 or NRG1:mScarlet3 co-expression with RBCS1A-GCaMP6s compared to the control, EV:mScarlet3 (NRG1^Δ20^:mScarlet3 – Wilcoxon, W=3446, p-value=0.208 and NRG1:mScarlet3 – Wilcoxon, W=5511, p-value<0.005). **(C)** Confocal micrographs of *nrg1/adr1* KO *N. benthamiana* leaf epidermal cells transiently expressing RBCS1A-GCaMP6s:GFP, RCaMP1h and NRG1:mTurqoise2 with and without XopQ. All images shown are single-plane images. Scale bars represent 10 μm. Raw fluorescence intensity data can be found at Data S2. Imaging was performed at 26-28-hours post-agroinfiltration using a minimum of two-leaf patches per sample. (**D**) Box and dot plot showing the RCaMP1h:GCaMP6s relative intensity per chloroplast for 80 chloroplasts from two separate data collections. Each chloroplast is highlighted with a turquoise dot. The mean value for each condition is shown as a black dot. Asterisks indicate statistically significant difference of XopQ co-expression with NRG1:mTurqoise2 compared to the GUS control (XopQ – Wilcoxon, W=2, p-value<0.005). (**E**) Confocal micrographs of *N. benthamiana* wild-type leaf epidermal cells transiently expressing NRG1:mScarlet3 and RBCS1A-GCaMP6s:GFP. Images shown are single-plane images from a time series collected over an hour time-period 27-28 hours post agroinfiltration. Dashed lines in overview correspond to inset. Lines (a to b) in the insets correspond to line intensity plots depicting the relative fluorescence across the marked area. A second movie from a separate data collection can be found at Fig. S11.

To elucidate the temporal dynamics of NbNRG1-dependent depletion of Ca^2+^ from the chloroplast, we tracked stromal Ca^2+^ sensor (RBCS1A-GCaMP6s) fluorescence over time on various chloroplasts in cells labelled by NbNRG1:mScarlet3. Time-lapse imaging over 60 minutes linked NbNRG1 activation directly to Ca^2+^ export. As NbNRG1:mScarlet3 shifted from a diffuse cytoplasmic localization to punctate signal surrounding chloroplasts, stromal GCaMP6s fluorescence declined in the same chloroplasts (Fig. 2E and Fig. S11). Subsequently, with the onset of cell death an overall reduction in the NbNRG1:mScarlet3 is observed. This indicates that puncta formation at the chloroplast envelope directly corresponds to Ca^2+^ efflux from the stroma. Collectively, these data establish that, upon activation, NbNRG1 acts at the chloroplast envelope to mediate Ca^2+^ release from the chloroplast stroma into the cytosol, and that this channeling activity critically depends on the integrity of the most N-terminal region of the CC domain.

### CC_R_-NLRs possess an extended N-terminus relative to canonical CC-NLRs

The distinct membrane targeting and Ca^2+^ channeling activities of NbNRG1 at organellar membranes imply evolutionary divergence from the plasma membrane-centric mechanism of canonical CC-NLRs. To understand this functional specialization, we examined the sequence and structural evolution of CC_R_-NLR N-termini across land plants. To clarify the sequence diversity of the N-terminus of CC_R_-NLRs, we searched for N-terminal motifs in CC_R_-NLRs across land plants, using canonical CC-NLRs as an outgroup (Fig. S12 and Data S4 to S12). As previously reported, canonical CC-NLRs including ZAR1, Sr35 and the NRC clade members, share a conserved N-terminal MADA motif which is required for cell death function and is orientated towards the plasma membrane upon resistosome formation (*6, 7, 16, 41*). In contrast, CC_R_-NLRs showed several distinct N-terminal motifs across the land plants, with an initial divergence of two CC_R_-NLR groups in the monilophytes and the gymnosperms. Subsequently, ADR1- and NRG1-specific N-terminal motifs appear to have evolved during divergence of the angiosperms. This data suggests that, unlike canonical CC-NLRs, CC_R_-NLRs evolved multiple distinctive N-terminal motifs during the evolution of the land plants.

Given that NbNRG1 targets organelle membranes, we asked whether this feature is encoded in its unique N-terminal architecture, and whether this enables the NbNRG1 resistosome to function at membranes beyond the plasma membrane. To address this, we examined the structural divergence of the N-terminal helices of CC_R_-NLRs. Because no resolved structures of activated CCR-NLR resistosomes are available, we modeled CC_R_-NLR oligomers using AlphaFold 3 (AF3) and compared them to experimental structures of CC-NLR resistosomes (*14, 42*). AF3 predictions revealed that CC_R_-NLR resistosomes possess a markedly elongated CC funnel relative to canonical CC-NLRs (Fig. 3A, Fig. S13, Data S13). Notably, the NbNRG1 funnel extends ∼50 Å beyond those of ZAR1, Sr35, and NRC4, potentially exceeding the thickness of a single lipid bilayer (Fig. 3A) (*43*). Predictions of the resting state likewise reveal a uniquely extended fourth α-helix in NbNRG1 absent from canonical CC-NLRs (Fig. S14). We also used AF3 to evaluate the truncated variants employed in our studies (NbNRG1^Δ14^ and NbNRG1^Δ20^). Both formed intact CC domain structures similar to full-length NbNRG1 but with shorter N-terminal regions (Fig. S15A), reinforcing the functional importance of the extended CC domain (Fig. 2A and B, Fig. S5, and Fig. S10). Analysis of the electrostatic potential of the predicted NbNRG1 CC domain revealed, as observed for *A. thaliana* NRG1 (*25*), positive, negative, and neutral regions at the periphery, lumen, and outer surface, respectively (Fig. S15B and C). Overall, these in silico analyses indicate that NbNRG1 possesses an extended CC domain architecture relative to canonical CC-NLRs, which may underlie its ability to target diverse membrane compartments.

**Fig. 3.**
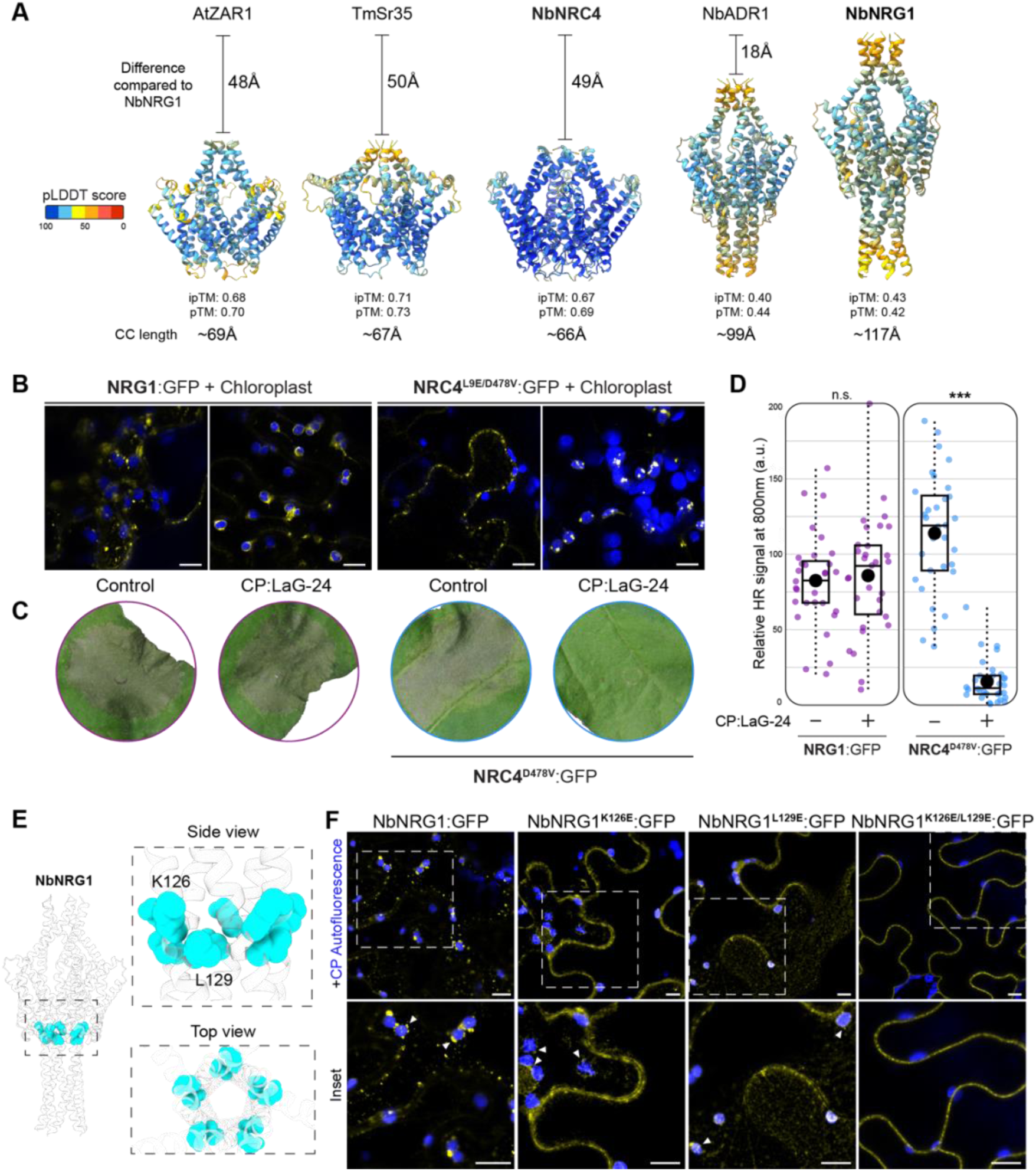
An extended coiled-coil domain architecture enables NbNRG1 organellar targeting. (**A**) AlphaFold3 predicted coiled-coil (CC) domain models of AtZAR1, TmSr35, NbNRC4, NbADR1 and NbNRG1. Structures are colored by pLDDT values. NbNRG1 and NbADR1 are modelled as pentamers as we observed higher pTM and ipTM scores compared to hexamers. The difference in angstroms to the NRG1 CC domain is shown for each CC modelled. The total length of each CC domain can be found below the respective predicted structure. AlphaFold3 model confidence analysis can be found at Fig. S13. (**B**) Confocal micrograph of *N. benthamiana* leaf epidermal cells transiently expressing NbNRG1:GFP or NRC4^L9E/D478V^:GFP with either a control or NtHR:LaG-24 (CP:LaG-24). Chloroplasts are visualized using auto-fluorescence at 648-709 nm. All images shown are single-plane images. Scale bars represent 10 μm. Imaging was done with at least two separate leaf patches from different plants. (**C**) Representative cropped leaf images from wild-type *N. benthamiana* plants showing HR after expression of NRG1:GFP and NRC4^D478V^:GFP either with CP:LaG-24 nanobody or control. (**D**) Box and dot plot showing the normalized HR signal at 800 nm (infrared) for 31 leaves, from two separate technical repeats. For (C) and (D), each leaf is highlighted with purple for NRG1:GFP and blue for NRC4^D478V^:GFP. For (D), the mean value for each condition is shown as a black dot. Asterisks indicate statistically significant difference of CP:LaG-24 treatment compared to the control (NRG1:GFP – t test, p-value=0.7089 and NRC4^D478V^:GFP – Wilcoxon, W=7, p-value<0.005). Raw HR intensity can be found at Data S3. **(E)** AF3 model for NbNRG1 highlighting the residues studied, K126 and L129, colored light blue. **(F)** Confocal micrographs of *nrg1/adr1 N. benthamiana* leaf epidermal cells transiently expressing NbNRG1:GFP, NbNRG1^K126E^:GFP, NbNRG1^L129E^:GFP or NbNRG1^K126E/L129E^:GFP. Chloroplasts are visualized using auto-fluorescence at 648-709 nm. Images shown are single-plane images. Dashed lines in top panels correspond to insets (bottom panel). Imaging was done with at least two separate leaf patches per panel.

### Unlike NbNRG1, enrichment of the canonical CC-NLR NbNRC4 at the chloroplast abolishes its activity

Ca^2+^ channeling and downstream HR functions are conserved amongst canonical CC-NLRs and CC_R_-NLRs, yet their CC domains are predicted to differ sharply in length (Fig 3A and Fig. S14). As the elongated CC_R_ domain of NbNRG1 is essential for Ca^2+^ efflux from chloroplasts (Fig. 2A and B, Fig. S9, Fig. S11 and Fig. S15), we asked whether the shorter CC domain of the CC-NLR helper NbNRC4 could perform the same function. To address this, we redirected the auto-active NbNRC4^D478V^:GFP and full-length NbNRG1:GFP to the chloroplast surface by co-expression with a GFP-binding nanobody anchored in the chloroplast outer envelope by fusion with the N-terminus of CHUP1 (CP:LaG-24) (Fig. 3B to D) (*44, 45*). Nanobody trapping at the chloroplast did not significantly alter the soluble levels of either NbNRG1:GFP, NbNRC4^D478V^:GFP or GFP control, although protein levels for NbNRC4^D478V^ with or without CP:LaG-24 were lower compared to NbNRG1 (Fig. S16). Chloroplast trapping abolished the HR function of NbNRC4^D478V^:GFP but not that of NbNRG1:GFP (Fig. 3B to D). Our stromal Ca^2+^ measurements with the red fluorescent sensor RBCS1A-jRGECO1a mirrored these phenotypes (*39, 46*). NbNRG1 trapped at the chloroplast envelope depleted stromal Ca^2+^ stores but NbNRC4^D478V^ and the GFP control did not, showing significantly higher sensor:GFP ratios than NbNRG1 (Fig. S17). As a control, we trapped NbNRG1:GFP and NbNRC4^D478V^:GFP at the plasma membrane using the same anti-GFP nanobody (Lti6b:LaG-24 or PIP2a:LaG-24) (*45, 47*). Trapping did not affect HR activity (Fig. S18), confirming that cell-death function is intact. For NbNRG1, although a fraction of protein escaped to its native location, residual puncta show that some were still retained at the plasma membrane. Together with our structural modeling, these results indicate that the distinct CC_R_ domain architecture of divergent helper NLRs allows them to function at specific membrane interfaces.

### The extended fourth α-helix underpins NbNRG1 membrane targeting and HR activities

Compared to canonical CC-NLRs, NbNRG1 contains a uniquely extended fourth α-helix (Fig. S14), previously shown to be essential for HR function in *Arabidopsis* NRG1 variants (*6, 25*). To determine how this region contributes to organellar targeting, we generated single and double point mutants within this extended helix (Fig. 3, E and F, and Fig. S19). Single mutants, NbNRG1^K126E^:GFP and NbNRG1^L129E^:GFP, exhibited partial organelle and plasma membrane localization and retained HR activity similar to that of wild-type NbNRG1 (Fig. 3, E and F, and Fig. S19), consistent with the incomplete plasma membrane trapping observed previously with Lti6b:LaG-24 and PIP2a:LaG-24 nanobodies (Fig. S18). In contrast, the double mutant NbNRG1^K126E/L129E^:GFP was fully retained at the plasma membrane and showed a marked reduction in HR (Fig. 3, E and F, and Fig. S19), indicating that exclusive plasma membrane localization impairs NbNRG1’s cell-death execution function. Both K126 and L129 face the outside of the NbNRG1 CC funnel and do not alter the predicted pore architecture (Fig. 3E), consistent with NbNRG1^K126E/L129E^:GFP retaining HR activity relative to the pore-deficient NbNRG1^Δ20^:GFP control (Fig. S19, A and B). These residues are also distal to the EDS1-SAG101 interaction surface on NRG1 (*22–23*), suggesting that upstream activation is likely preserved and that the observed phenotype primarily reflects altered membrane targeting. Protein levels mirrored these phenotypes. NbNRG1^K126E/L129E^:GFP accumulated to higher levels than the single mutants and wild-type NbNRG1, consistent with reduced cell death (Fig. S19C). Together, these data demonstrate that the extended fourth α-helix is the critical structural element that directs NbNRG1 to the correct organellar membrane environment and is essential for its HR execution function.

### Chloroplast targeting is a conserved feature amongst plant CC_R_-NLRs

Given the evolution of unique N-terminal motifs in CC_R_-NLRs across the land plants, we wondered whether chloroplast targeting is a conserved feature of CC_R_-NLRs. To investigate the conservation of chloroplast targeting, we examined the sub-cellular localisation of 5 CC_R_-NLRs that span ∼360 million years of land-plant divergence (Fig. 4A). We selected 4 NRG1 subclass members from across the angiosperms, as well as one from the monilophytes pre-dating the divergence of the NRG1 and ADR1 subclasses, which are the only CC_R_-NLR subclasses found in flowering plants (Fig. S12). Remarkably, all 5 CC_R_-NLRs accumulated as discrete puncta on the chloroplast outer membrane (Fig. 4B) and like NbNRG1, had extended CC domain structures compared to the canonical CC-NLR NbNRC4 (Fig. S20). These observations extend our findings beyond the model species *N. benthamiana* and suggest that chloroplast targeting is a conserved feature of NRG1-like CC_R_-NLRs, possibly predating the divergence of monilophytes and seed plants over 360 million years ago (*48*).

**Fig. 4.**
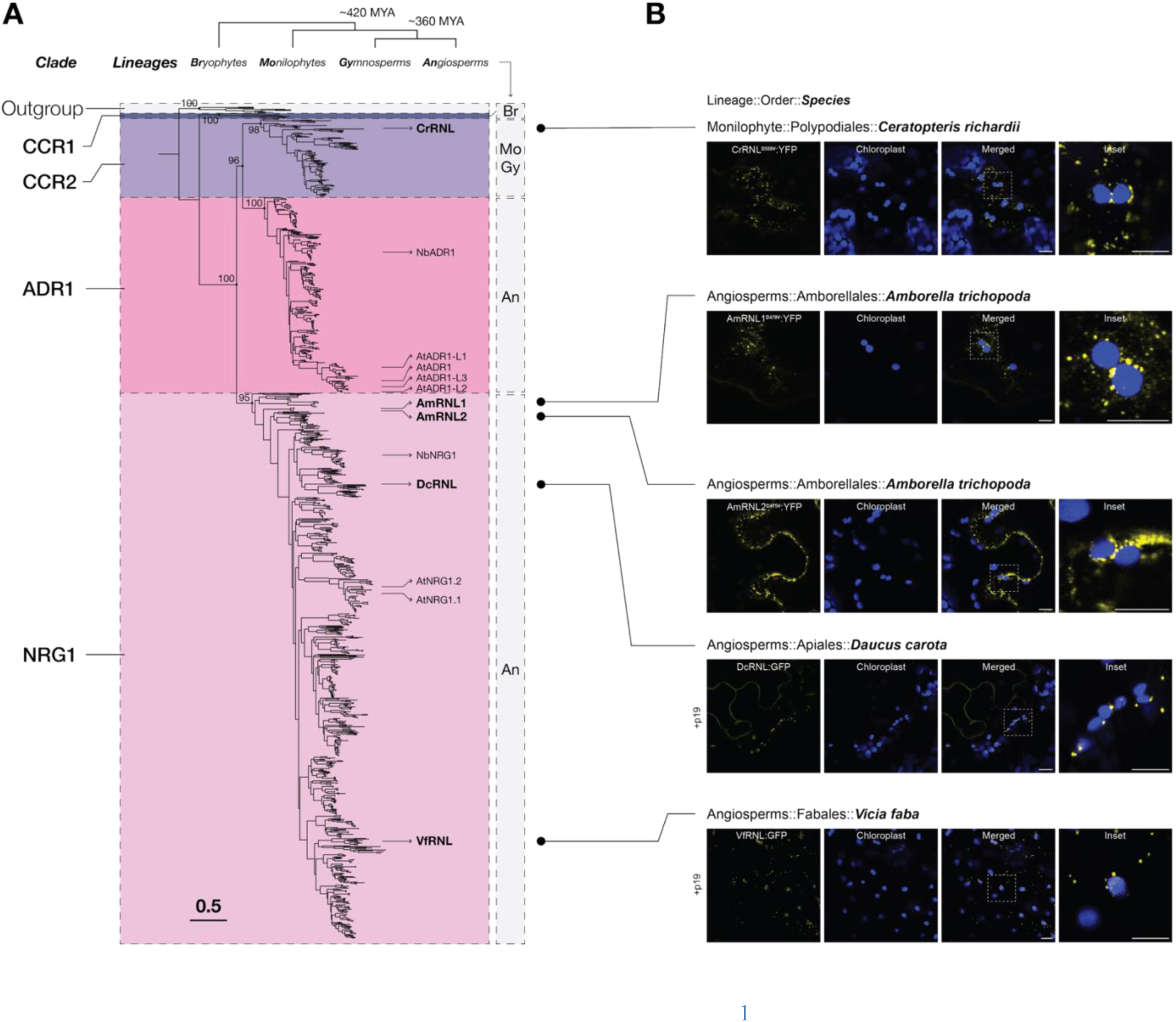
Chloroplast targeting is a conserved feature amongst CC_R_-NLRs (**A**) The phylogenetic tree illustrates the relationships among 56,280 CCR-NLRs based on their NB-ARC domain sequences. Four well-supported clades are highlighted: the CCR1 and CCR2 clades (purple), the ADR1 clade (pink) and the NRG1 clade (lilac). The tree is rooted with sequences from AtZAR1, NbZAR1, and NRCs from tomato as the outgroup. (**B**) Confocal micrographs of *N. benthamiana* leaf epidermal cells transiently expressing C-terminally YFP or GFP fused NRG1-like CC_R_-NLRs identified from the phylogenetic analysis in (A). Silencing suppressor p19 was co-expressed with wild-type DcCC_R_-NLR and VfCC_R_-NLR. For CrCC_R_-NLR and AmCC_R_-NLR1/2 auto-active D to V mutants were coupled with L to E α-helix1 mutations to allow imaging. Chloroplasts are visualized using auto-fluorescence at 648-709 nm. All images shown are single-plane images. Dashed lines in merged panels correspond to insets. Scale bars represent 10 μm.

## Discussion

CC_R_-NLRs constitute one of the oldest NLR clades, persisting from mosses to angiosperms (*9*). Here, we show that the Solanaceae CC_R_-NLR NbNRG1, upon activation, undergoes a striking shift from a diffuse cytosolic pool to puncta on internal organelle membranes (Fig. 1). NbNRG1 targets the chloroplast envelope and facilitates efflux of stromal Ca^2+^ into the cytosol, showing that NLRs do not exclusively rely on extracellular calcium reservoirs for immune signalling (Fig. 2 and Fig. S9). We show that organelle targeting is conserved amongst NRG1 orthologues across the angiosperms and may even date back to over 360 million years, since a monilophyte CC_R_-NLR predating the evolution of the ADR1 and NRG1 clades also localises to this membrane (Fig. 4). This organelle targeting starkly contrasts with the localisation of the canonical CC-NLR NRC4, which exclusively targets the plasma membrane (Fig. S4), suggesting a unique mechanism for cell death activation beyond plasma membrane pore formation and cation channel activity (*8, 18, 19, 26*).

Despite the overall conservation of the CC domain four-helix bundle, the N-terminal sequence of plant CC-NLRs is surprisingly diverse (Figs. S12 and S13) (*17, 28*). However, the biological drivers of this diversification remain unknown. Our structural models of the NbNRG1-like resistosomes predict an extended N-terminal funnel-shaped structure compared to that of canonical CC-NLRs, a unique feature of CC_R_-NLRs that could enable these proteins to span double-membrane organelles, such as chloroplasts (Fig. 1, F to H; Fig. 3A; Fig. S15) (*43*). NbNRG1 exhibits an extended fourth α-helix compared to canonical CC-NLRs (Fig. S14), and point mutations within this helix drastically altered organelle membrane localization, diverting NbNRG1 to the plasma membrane where its HR function was hampered (Fig. 3, E and F, and Fig. S19). The mutated residues face the outer-face of the CC funnel (Fig. 3E), indicating a potential role in membrane docking and interactions, rather than a direct involvement in the channeling of cations, as the mutants exhibited cell death compared to our negative control. Together, these findings suggest that while the four-helix bundle architecture of CC domains is conserved, the extended fourth α-helix in NbNRG1 confers functional diversification by enabling distinct membrane targeting capabilities.

In this context, it is notable that *Arabidopsis* NRG1 variants also carry an extended helix-4, yet previous studies have so far reported primarily plasma-membrane-localized resistosomes (*6, 25*). These analyses were largely performed with autoactive AtNRG1 alleles expressed in *N. benthamiana* rather than with native *Arabidopsis* NRG1 activated in its endogenous cellular environment. Experiments were not necessarily designed to systematically probe potential organelle localization. Our mutational data show that modest changes in helix-4 residues are sufficient to switch NbNRG1 between chloroplast and plasma-membrane targeting. A full exploration of whether *Arabidopsis* NRG1 can also access organellar membranes will require further analysis.

The ability of NRG1 to target calcium-rich organelles such as chloroplasts, mitochondria, and the ER may enable it to tap into alternative intracellular calcium stores, diversifying the cell death immune response (*49*). Using multiple Ca^2+^ sensors for the chloroplast stroma and the cytoplasm, we demonstrate that, upon activation by the bacterial effector XopQ, the unique localization pattern of NbNRG1 at the chloroplast envelope is indeed followed by Ca^2+^ efflux, and that the extended length of the NbNRG1 CC domain is necessary for cation channeling from the chloroplast (Fig. 2). By diversifying its subcellular localization, NRG1 may have evolved to initiate immune responses via multiple membrane interfaces. This could represent a multi-compartmental defense strategy to counteract pathogen effectors that suppress activated NLR functions at the plasma membrane and therefore sustain robust immunity.

Nanobody trapping showed that NbNRG1, but not NbNRC4, can facilitate Ca^2+^ efflux and HR when immobilized at the chloroplast envelope. Plasma membrane nanobody-trapped NbNRG1 and NbNRC4^D478V^ remained largely at the plasma membrane, although trapping was incomplete, with residual NbNRG1 puncta visible in the cytosol (Fig. S17 and Fig. S18). This partial retention parallels the behaviour of single NbNRG1 point mutants (K126E or L129E) in the extended fourth α-helix, which exhibited partial plasma membrane localization and retained HR activity comparable to wild-type NbNRG1 (Fig. 3, E and F, and Fig. S19). By contrast, the double mutant NbNRG1^K126E/L129E^, which fully localized to the plasma membrane, displayed reduced but not abolished HR, consistent with the positioning of the residues on the outer-surface of the funnel rather than the inner cation channel (Fig. 3E). These findings indicate that incomplete plasma membrane trapping, whether via nanobody tethering or single point mutation, permits sufficient organellar targeting to sustain HR, whereas full retention at the plasma membrane reduces this function. We propose that NbNRG1’s extended N-terminus and elongated CC_R_ domain architecture broadens membrane compatibility, by enabling tethering and/or stabilizing pore formation at the chloroplast outer envelope, while NRC4 likely depends on plasma membrane cofactors or is constrained by membrane thickness, curvature and lipid composition. For example, the thickness of the chloroplast outer membrane is between 50-60 Å (*43*), which is the difference we measured between the CC domain structures of NbNRG1 and NbNRC4 (Fig. 3A). These results highlight subcellular localisation and membrane context as decisive determinants of NLR function.

The localisation of NbNRG1 to organelles supports a model in which NLR-mediated immune signaling is more compartmentalized and complex than anticipated (Fig. S21). Our findings align with recent research emphasizing the critical role of chloroplasts in immunity (*50, 51*). Upon immune activation, chloroplasts downregulate photosynthesis and produce reactive oxygen species (ROS), defense hormones, and facilitate signaling through the MAPK pathway (*49–56*), processes that are tightly linked to NLR-mediated immunity. We observed NRG1 puncta at the stromules (Fig. S6), which act as conduits to transport pro-defense signals, like ROS (*35*). In addition to mediating Ca^2+^ efflux, activated NRG1 may interact with these chloroplast immune processes, further linking organellar function to NLR-triggered immunity.

Our findings reveal hidden functional diversity among CC domain NLRs, with sequence features like N-terminal extensions tuning membrane preference and channel activity. Decoding this permissive-membrane tethering code could enable engineering of membrane-tuned NLRs for more versatile and durable plant immunity.

## Acknowledgments

We thank the Imperial College FILM facility for their technical expertise and provision of our microscopy equipment.

## Funding

Biotechnology and Biological Sciences Research Council (BBSRC) Impact Acceleration Fund BB/X511055/1 (T.I.), BBSRC BB/X016382/1 (E.L.H.Y. and T.O.B.)

National Science and Technology Council NSTC-112-2628-B-001-007, NSTC 113-2927-I-001-514 (H.Y.W. and C.H.W.)

The Gatsby Charitable foundation (A.T., S.K., B.C. and J.D.G.J.) BBSRC BB/P012574 (Plant Health ISP) (A.T. and S.K.) BBSRC BBS/E/J/000PR9795 (A.T. and S.K.), BBSRC BBS/E/J/000PR9796 (Plant Health ISP – Response) (A.T. and S.K.) BBSRC BBS/E/J/000PR9797 (Plant Health ISP – Susceptibility) (A.T. and S.K.) BBSRC BBS/E/J/000PR9798 (Plant Health ISP – Evolution) (A.T. and S.K.) BBSRC BB/V002937/1 (A.T. and S.K.), BBSRC BB/Y514201/1 (J.K.)

UK Research and Innovation (UKRI) EP/Y032187/1 (A.T. and S.K.) European Research Council (ERC) BLASTOFF 743165 (A.T. and S.K.) BBSRC BB/X010996/1 (Advancing Plant Health) (K.S.C. and P.C.)

## Author contributions

Conceptualization: T.I., F.J.K., J.K., T.O.B.

Methodology: T.I., F.J.K., A.T., J.K., T.O.B.

Investigation: T.I., F.J.K, L.W., S.J., E.L.H.Y., H.Y.W., A.T., J.K., C.V., N.E., V.A., K.S.C., B.C., T.O.B.

Visualization: T.I., E.L.H.Y., A.T., T.O.B.

Funding acquisition: T.O.B., C.H.W., S.K., J.D.G.J., P.C.

Project administration: T.I., F.J.K., J.K., T.O.B. Supervision: T.I., T.O.B., C.H.W., S.K., J.D.G.J., P.C.

Writing – original draft: T.I., F.J.K., T.O.B

Writing – review & editing: T.I., F.J.K., E.L.H.Y., A.T., J.K., S.K., J.D.G.J., C.H.W., P.C., T.O.B.

## Competing interests

T.O.B. and S.K. receive funding from industry and co-founded a start-up company (Resurrect Bio Ltd.) on NLR biology. S.K. and J.K. filed patents on NLR biology.

## Data and materials availability

Constructs generated for this study will be subjected to material transfer agreements (MTAs) and made available upon request. All data are available in the main text or the supplementary materials.

## Supplementary Materials

### Materials and Methods

Figs. S1 to S21

Tables S1 to S2

Data S1 to S13

## Notes

https://doi.org/10.5281/zenodo.18593781

### Competing Interest Statement

T.O.B. and S.K. receive funding from industry and co-founded a start-up company (Resurrect Bio Ltd.) on NLR biology. S.K. filed patents on NLR biology.

### Summary of Updates

The updated version shifts the emphasis to a chloroplast-envelope-centred model. It adds functional experiments that directly link chloroplast association to immune output, and it introduces calcium-signalling measurements consistent with chloroplast-to-cytosol calcium mobilisation. It also includes nanobody-relocalization/tethering approaches to test membrane specificity, showing NRG1 remains functional when driven to chloroplast membranes while canonical CC-helper NRC4 does not. Finally, it adds tighter structure-function mapping by pinpointing a more specific determinant with localization and activity mutants.

