## Supplementary Materials for "A helper NLR channels organellar calcium to trigger plant immunity"

**The PDF file includes:**

Materials and Methods
Figs. S1 to S21
Tables S1 to S2

**Other Supplementary Materials for this manuscript include the following:**

Data S1 to S13

### Materials and Methods

#### Molecular cloning

We employed Golden Gate Modular Cloning (MoClo) kit *(57)* along with the MoClo Plant Parts kit *(58)* for cloning purposes. Remaining clones were constructed using Gibson assembly of PCR products with 30 bp homology overhangs into the binary vector pK7WGF2, as previously described *(59)*. All clones were sequence verified before transformation into *Agrobacterium tumefaciens* GV3101/pMP90. Cloning design and sequence analysis were performed using Geneious Prime (v2023.2.1; www.geneious.com). Detailed information on plasmid construction can be found in Table S1 and the sequences of oligos used for cloning can be found in Table S2.

#### Plant material

Wildtype and transgenic *Nicotiana benthamiana* plants were grown in a regulated growth chamber set at 24°C. The transgenic *nrg1/adr1* double KO plants were obtained from the Schattat lab *(60)*. The *epss* knockout plants were provided by the Parker lab *(61)*. To generate *nrg1* KO plants, the following primer pairs were used to manufacture guide RNA sequences targeting *nrg1*: CAGTATTCGATGACATCGAG(AGG) and GCTTGAGGAGAAAGAGAAGG(TGG). The primers used for genotyping *nrg1* KO plants were GAGCTGCTCTTGGTCCAGTT and TTGATATCATCCTCTTGACAAAGC. The cultivation medium of the plants consisted of a blend of organic soil, mixed in a 3:1 ratio of Levington's F2 soil with sand and Sinclair's 2-5 mm vermiculite. The plants were exposed to intense light and maintained under long-day conditions, with a photoperiod of 16 hours of light and 8 hours of darkness. The experiments were conducted on plants aged between 4 and 5 weeks.

#### *Agrobacterium*-mediated transient gene expression in *N. benthamiana*

*Agrobacterium*-mediated transient gene expression was carried out via agroinfiltration as described previously *(62)*. *A. tumefaciens* GV3101/pMP90 strains containing binary plasmids were grown on LB plates or in liquid LB media containing the appropriate antibiotics (kanamycin 100 µg/mL or spectinomycin 100 µg/mL) including gentamycin (50 µg/mL). The cultures were collected and first washed with water and then resuspended in an agroinfiltration buffer composed of 10 mM MES and 10 mM $MgCl_2$ (pH 5.7). The $OD_{600}$ of the bacterial suspension was measured and then adjusted according to the specific construct and experimental requirements. Finally, the prepared bacterial suspension was infiltrated into 4 to 5-week-old *N. benthamiana* leaves using a 1 ml Plastipak syringe without a needle.

#### Chloroplast isolation

Deveined leaf tissue (15g) from *N. benthamiana* agroinfiltrated 2-days prior was homogenized in ice-cold grinding buffer (330 mM sorbitol, 10 mM $Na_4P_2O_7$, 2.5 mM EDTA, 40 mM D-isoascorbate, 5 mM $MgCl_2 \cdot 6H_2O$, pH 6.5). Homogenate was filtered through cheesecloth and centrifuged (4000 × g, 3 min, 4 °C). The pellet was resuspended in 100 µL resuspension buffer (330 mM sorbitol, 5 mM $MgCl_2 \cdot 6H_2O$, 10 mM KCl, 2.5 mM EDTA, 50 mM HEPES, pH 7.6). A 50% (v/v) Percoll solution was prepared in double-strength resuspension buffer (660 mM sorbitol,

5 mM $MgCl_2 \cdot 6H_2O$, 10 mM KCl, 2.5 mM EDTA, 50 mM HEPES, pH 7.6), and 6 mL was added to the resuspended pellet. Samples were centrifuged (4000 × g, 7 min, 4 °C), and the intact chloroplast fraction was collected, resuspended in 100 µL resuspension buffer, and stored on ice for confocal microscopy.

**Confocal laser scanning microscopy**

Confocal microscopy was performed 1-3 days post-agroinfiltration as described previously *(62)*. Leaf tissue samples were collected using a size 4 cork borer, submerged in distilled water, and mounted live on glass slides for imaging. The abaxial side of the leaf tissue was imaged using a Leica STELLARIS 5 inverted confocal microscope with a 63x water immersion objective lens. The emission wavelengths for GFP (mGOLD/StayGold), RFP (mScarlet3) and BFP (mTurqoise2) tags were set to 495-550 nm, 570-620 nm and 402-457 nm, respectively. The confocal images were analyzed using ImageJ.

**Image processing and data analysis**

Confocal microscopy images were processed using Leica LAS X software and ImageJ. Image analysis and quantification for cell death experiments were conducted with ImageJ. Data were represented using box-and-dot plots created with ggplot2 in R *(63)*.

**Cell death assay**

Cell death elicitors were introduced into the abaxial side of *N. benthamiana* leaves via agroinfiltration. At 3 dpi, the leaves were detached and subjected to both daylight and infrared imaging. Infrared imaging was performed using the 800 nm channel of the Bio-Rad ChemiDoc MP Imaging System. The resulting infrared images were analyzed with the "Volume Analysis" function in Bio-Rad Image Lab software. The HR signal was determined by dividing the signal intensity of the cell death spot by its area and then normalizing the value on a scale of up to 10 points. The normalized HR signal was plotted using ggplot in R *(63)*.

**Blue native PAGE and immunoblot analysis**

Proteins of interest were expressed in *N. benthamiana* leaves by agroinfiltration and leaf samples were taken at 1 dpi for Western blot and 2 dpi for blue native PAGE. For each sample, six leaf discs were taken using a size 4 cork borer and proteins were extracted, resolved and visualised as previously described *(62)*. For blue native PAGE analysis, proteins were extracted without denaturation, resolved and visualised using protocols which have been previously described *(64)*. For detection of GFP-tagged proteins, polyclonal anti-GFP antibodies produced in rabbit were used as primary antibodies (PABG1, ChromoTek) and anti-rabbit antibodies for horseradish peroxidase (HRP) detection (A9169, Sigma-Aldrich) were used as secondary antibodies. For detection of myc-tagged proteins, anti-myc antibodies produced in mouse (A00704, GenScript) were used as primary antibodies and anti-mouse antibodies for HRP detection (115-035-003, Jackson) were used as secondary antibodies.

**Structural analyses (AlphaFold3)**

The AlphaFold 3 web server (https://golgi.sandbox.google.com/) was used to model pentamers for AtZAR1, TmSr35, NbADR1 and NbNRG1, and hexamer for NbNRC2 with 50 oleic acids as a proxy for the plasma membrane (Data S13) *(42)*. AtZAR1 and TmSr35 were modeled as pentamers and NbNRC2 as a hexamer based on their cryoEM structures (PDB ID:6J5T *(7)*, 7XC2 *(16)* and 9FP6 *(20)*, respectively). The default seed was set to 1. From each modeling run, only the top-ranked model (model_0) was retained and processed. The ipTM and pTM values were extracted from the AlphaFold 3 JSON files using custom scripts. PAE plots for models were downloaded from the AlphaFold 3 web server. All structural predictions and their associated data are available from https://doi.org/10.5281/zenodo.18593781.

**Phylogenetics and motif analyses**

We extracted 110,819 NLR proteins from the NLRtracker outputs of 180 RefSeq plant proteomes *(65)*, available on Zenodo (Data S5) *(66)*. From these, 98,733 NLRs were identified based on specific domain architectures: "CNL", "CNLO", "CN", "OCNL", "CONL", "NL", "NLO", "ONL", "BCNL", "BNL", "BCN", "BCCNL", "BNLO", "BOCNL", "RNL", "TN", "TNL", "TNLO", and "TNLJ". To reduce redundancy, we deduplicated the dataset, retaining 63,125 unique NLR sequences. Further refinement involved filtering the sequences based on NB-ARC domain length, specifically retaining those between 250 and 400 amino acids. This step resulted in 56,280 NLRs for downstream analysis (Data S6 to S7). These NB-ARC domain sequences were aligned to the RefPlantNLR database using FAMSA v2.2.2 *(67, 68)* followed by phylogenetic tree construction using FastTree 2 v2.1.10 with the LG model *(69)*. From the resulting tree, we identified and extracted a $CC_R$-NLR clade comprising 1,140 sequences based on the inclusion of known reference ADR1 and NRG1 sequences. We then extracted 61 $CC_R$-NLR sequences from a collection of early diverging green plants and added to the extracted RefSeq sequences *(28)*. We then constructed a new NB-ARC tree with these $CC_R$-NLRs using MAFFT v7.525 with the [--anysymbol] option for alignment, ClipKIT v2.3.0 [-m gappy] for trimming the alignment, and IQ-TREE with options [-m JTT+F+I+G4 -B 1000 -T 8] *(70)*. AtZAR1, NbZAR1, SlNRC0, SlNRC1, SlNRC2, SlNRC3, SlNRC4a, SlNRC4b, and SlNRC6 sequences were used as outgroups *(66)*. The resulting tree was divided and annotated into four well-supported clades (Clades 1 to 4; as CCR1, CCR2, ADR1, NRG1, respectively). For the phylogenetic tree used in Fig. 4, the $CC_R$-NLR sequence from *Vicia faba* [Vf$CC_R$-NLR; Vfaba.Hedin2.R1.4g122640.1] was manually extracted and added to the compiled dataset *(71)*. A phylogenetic tree of the previous set and the added sequence was made similar to the previous step.

To identify conserved motifs within the $CC_R$ domains of these sequences, we extracted the domain sequences from the first residue to the start of the NB-ARC domain, excluding those shorter than 100 amino acids to avoid truncated sequences. MEME v5.5.7 [-maxw 30 -nmotifs 5 -mod zoops] was used to detect conserved motifs in the RPW8 domains of the four clades (Data S11) *(72)*. The N-terminal motifs identified by MEME were then used to create Hidden Markov Models (HMMs) using the hmmbuild software from the HMMER suite (Data S12). These HMMs were searched against the initial NLR datasets using hmmsearch with the [--max] option. Hits with e-values of 0.01 or lower were retained and mapped onto the $CC_R$-NLR phylogenetic tree. As a control, the MADA N-terminal motif was included to assess the distribution of N-terminal motifs *(41, 73)*. The resulting trees were visualized using iTOL *(74)*. All scripts used in this analysis are available on [https://github.com/amiralito/NRG1_Localization] and all NB-ARC amino acid sequences are available from https://doi.org/10.5281/zenodo.18593781.

### Supplementary Figures

**Fig. S1**

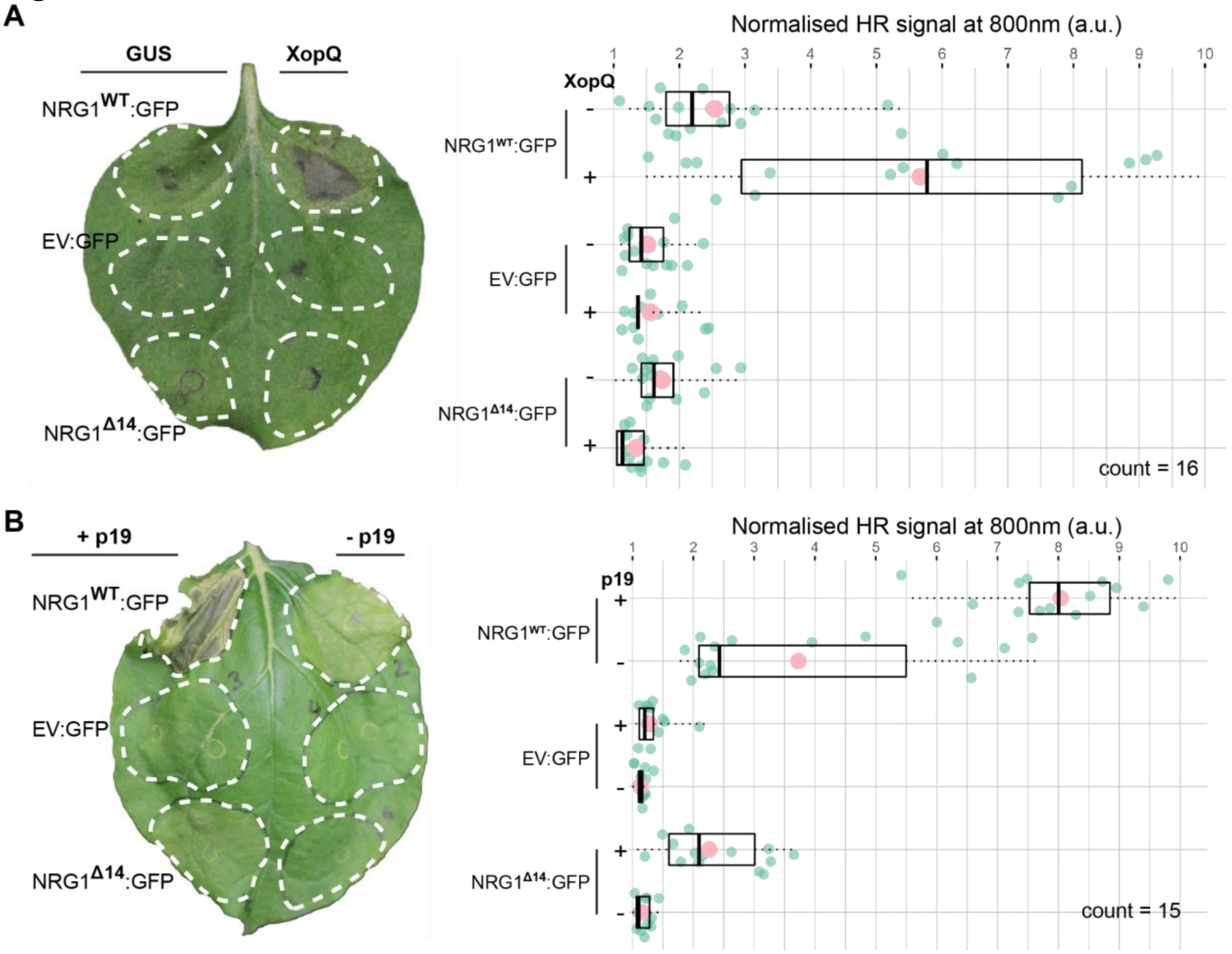


**Fig. S1 NbNRG1-dependent HR can be amplified by XopQ and p19**.

(**A**) A representative leaf picture from *N. benthamiana nrg1* KO plants showing HR after expression of NRG1:GFP, NRG1$^{\Delta14}$:GFP or EV:GFP with GUS control or XopQ. Box and dot plot showing the normalized HR signal at 800 nm (infrared) for 16 leaves over three replicates. Each leaf is highlighted with a green dot and the mean value for that condition in a larger pink dot. (**B**) A representative leaf picture from *N. benthamiana nrg1* KO plants showing HR after expression of NRG1:GFP, NRG1$^{\Delta14}$:GFP or EV:GFP with and without p19. Box and dot plot showing the normalized HR signal at 800 nm (infrared) for 15 leaves over two replicates. Each leaf is highlighted with a green dot and the mean value for that condition in a larger pink dot. Raw HR intensity can be found at Data S3.

**Fig. S2**

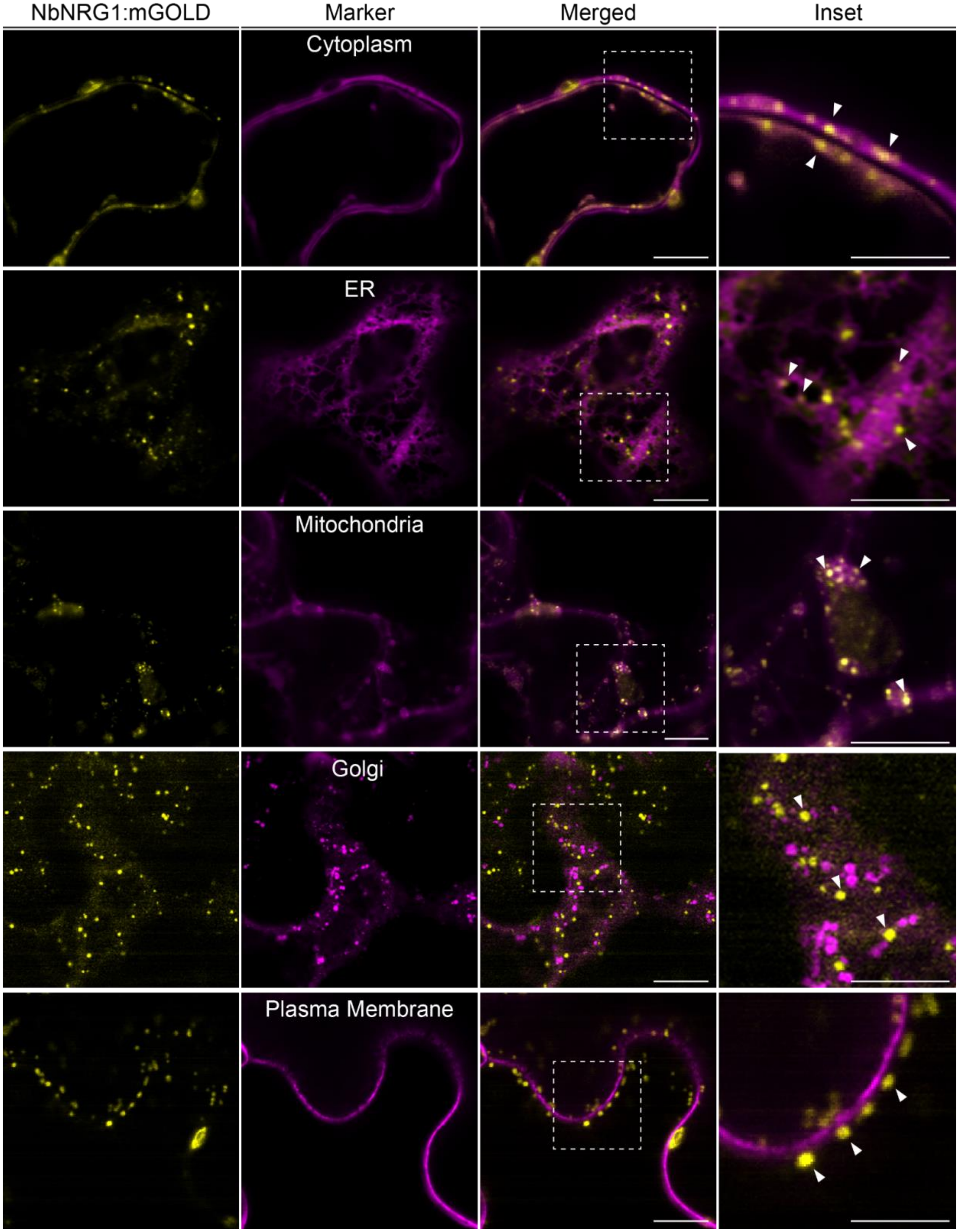
NbNRG1:mGOLD
Marker
Merged
Inset
Cytoplasm
ER
Mitochondria
Golgi
Plasma Membrane

**Fig. S2 Full-length NbNRG1 localizes to organellar membranes, like NRG1$^{\Delta 14}$.** Confocal micrographs of *N. benthamiana* leaf epidermal cells transiently expressing NbNRG1:mGOLD with SP:RFP:HDEL (ER), CTP1-RFP (mitochondria), GmMan11-49:mCherry (Golgi), RFP-Remorin1.3 (plasma membrane) or RFP:EV (Cytoplasm). Chloroplasts were visualized through autofluorescence. White arrows indicate NbNRG1:GFP puncta. Images shown are single-plane images. Dashed lines in the merged panel correspond to inset. Scale bars represent 10 µm. Imaging was done with at least two separate leaf patches per panel.

**Fig. S3.**

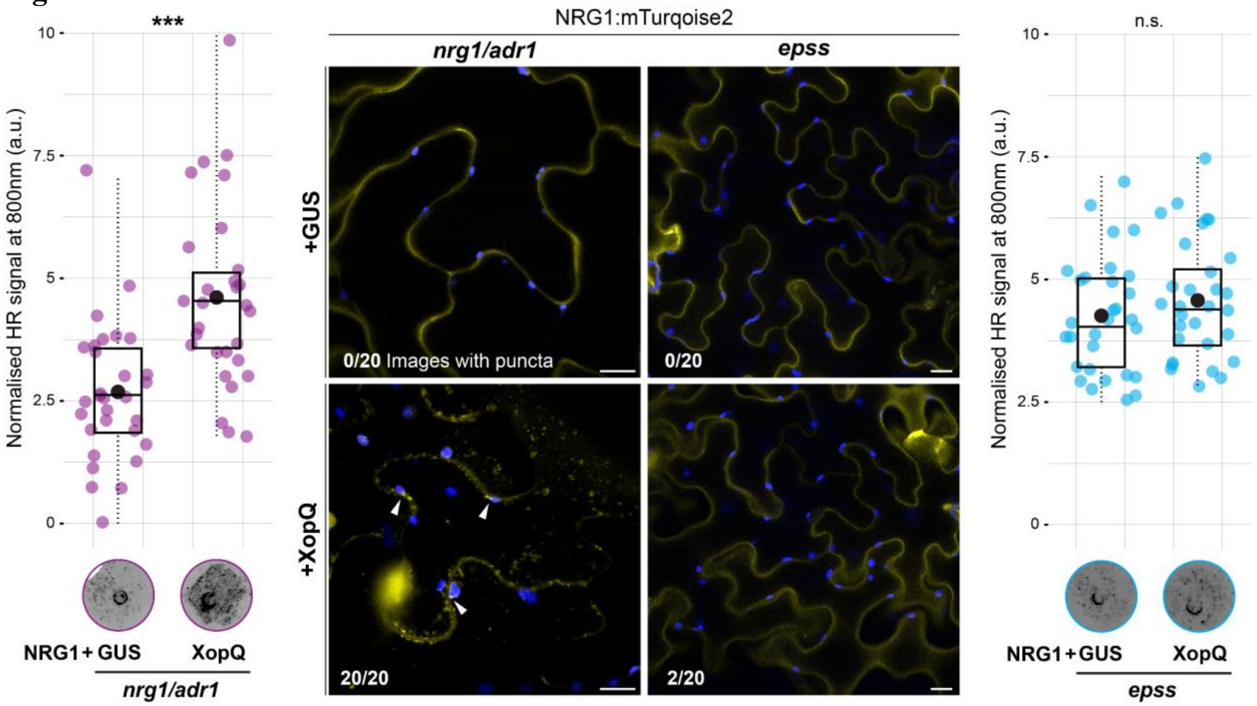


**Fig. S3 XopQ-dependent NbNRG1 activation at the chloroplast involves the canonical EDS1-SAG101 signalling module.**

Confocal micrographs (middle panels) and corresponding HR quantification (left and right panels) from *N. benthamiana nrg1/adr1* (purple) and *epss* (light blue) mutant plants transiently expressing NbNRG1:mTurqoise2 (p35S::NRG1:mTurquoise2 for microscopy and pAct2::NRG1:mTurquoise2 for HR assay) together with GUS or XopQ. Confocal imaging was performed at 24 hours post agroinfiltration, while HR was quantified from leaves imaged at 2 days post infiltration. In *nrg1/adr1* plants, XopQ co-expression induced prominent NbNRG1 puncta (arrowheads; 20/20 images) and significantly elevated HR compared to GUS controls (0/20 images with puncta; Welch two-sample t-test, $p=2.57 \times 10^{-5}$). In contrast, in *epss* plants, XopQ failed to induce NbNRG1 puncta (2/20 images) and did not significantly increase HR relative to GUS controls (Welch two-sample t-test, p=0.318). For the box and dot plots, each dot represents an individual leaf and black dots denote mean values. Statistical significance is indicated as *** ($p < 0.001$) or n.s. (not significant). Imaging was performed using at least two independent leaf patches from different plants. Scale bars, 10 µm.

**Fig. S4**

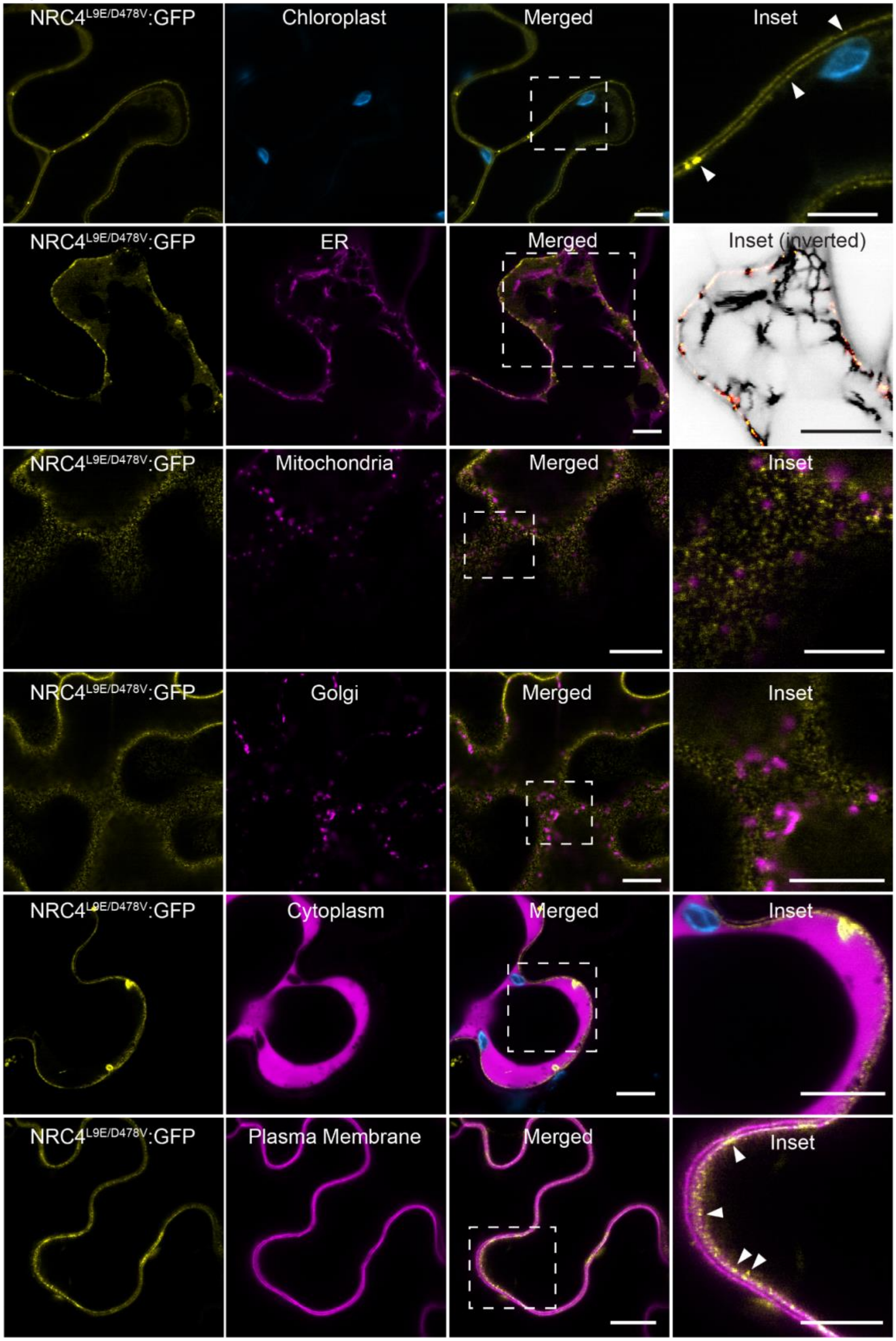


**Fig. S4 Auto-active CC-NLR NbNRC4 does not localize to organellar membranes.**

Confocal micrographs of *N. benthamiana* leaf epidermal cells transiently expressing NRC4$^{L9E/D478V}$:GFP with SP:RFP:HDEL (ER), CTP1-RFP (mitochondria), GmMan11-49:mCherry (Golgi), RFP-Remorin1.3 (plasma membrane) or RFP:EV (Cytoplasm). Chloroplasts were visualized through autofluorescence. White arrows indicate NRC4$^{L9E/D478V}$:GFP puncta. The ER, mitochondria, Golgi and cytoplasm panels were imaged at different Z-axis compared to plasma membrane and chloroplast panels, to be able to capture both the NRC4 punctate signal at the plasma membrane and respective organelle marker. Images shown are single-plane images. Dashed lines in the merged panel correspond to inset. Scale bars represent 10 µm. Imaging was done with at least two separate leaf patches per panel.

**Fig. S5**

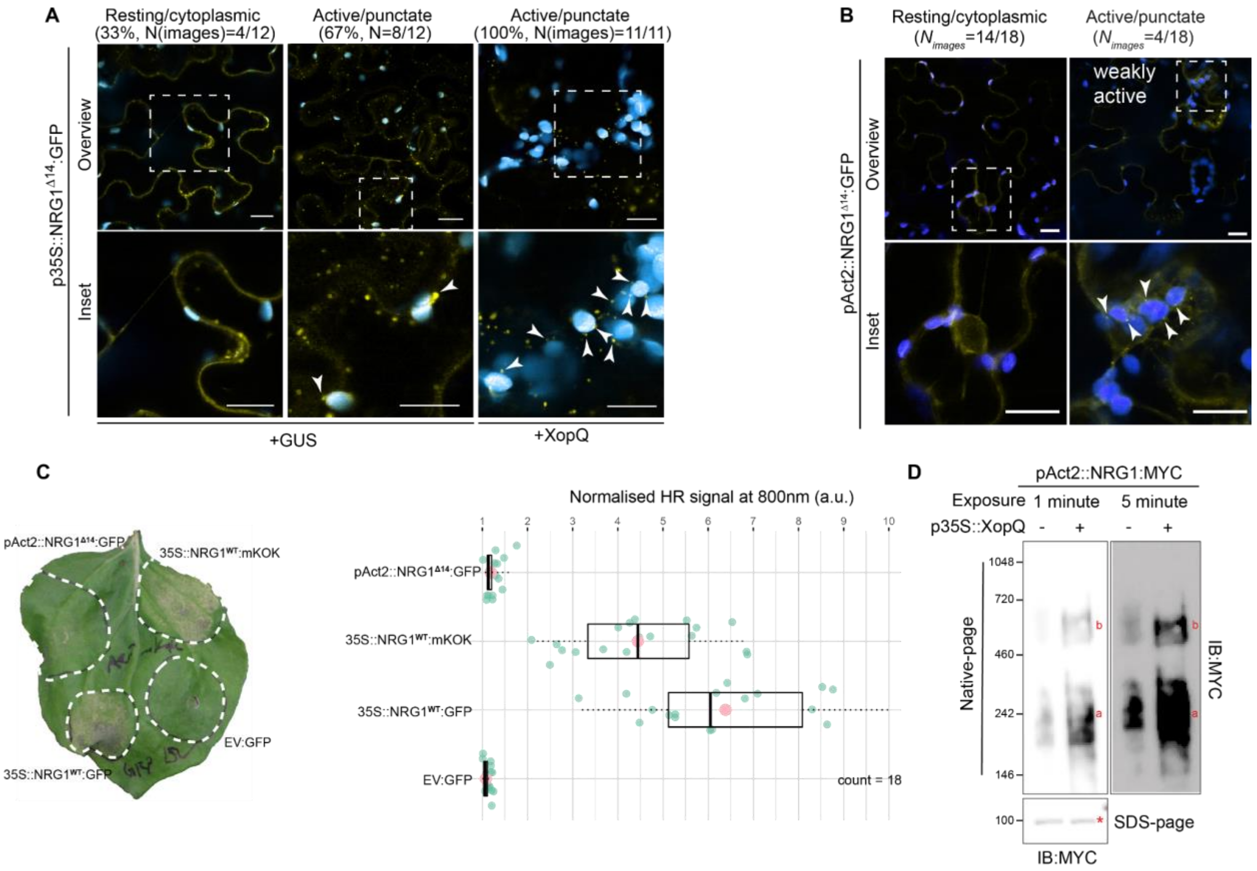


**Fig. S5 Activated NbNRG1$^{\Delta14}$ localizes to the chloroplast outer envelope.**
(**A**) Confocal micrographs of *N. benthamiana* leaf epidermal cells transiently expressing 35S::NRG1$^{\Delta14}$:GFP with XopQ or GUS control. Images shown are single-plane images. Dashed lines in overview correspond to inset. Arrows indicate examples of puncta around chloroplasts. Scale bars represent 10 µm. At least two different leaf patches were used for imaging. For GUS and XopQ treatments, 12 and 11 images were used, respectively. (**B**) Confocal micrographs of *N. benthamiana* leaf epidermal cells transiently expressing pAct2::NRG1$^{\Delta14}$:GFP. A total of 18 images were analysed from two-leaf patches. Images shown are single-plane images. Dashed lines in overview correspond to inset. Arrows indicate examples of puncta around chloroplasts. Scale bars represent 10 µm. Imaging was done with at least two separate leaf patches per panel. (**C**) A representative leaf picture from *N. benthamiana nrg1* KO plants showing HR after expression of pAct2:NRG1$^{\Delta14}$:GFP, 35S:NRG1$^{\Delta14}$:GFP or 35S:NRG1:mKOK. Box and dot plot showing the normalized HR signal at 800 nm (infrared) for 18 leaves. Each leaf is highlighted with a green dot and the mean value for that condition in a larger pink dot. Raw HR intensity can be found at Data S3. (**D**) Blue native PAGE analysis of full-length pAct2::NbNRG1:MYC activation by XopQ in *N. benthamiana nrg1/adr1* KO leaf tissue. SDS-PAGE analysis of reduced state of NbNRG1 as a control for soluble protein quantity. Letters a and b indicate two distinct species detected using MYC immunoblotting (a – resting and b – activated). Red asterisk indicates expected band size for soluble protein in SDS-PAGE.

**Fig. S6**

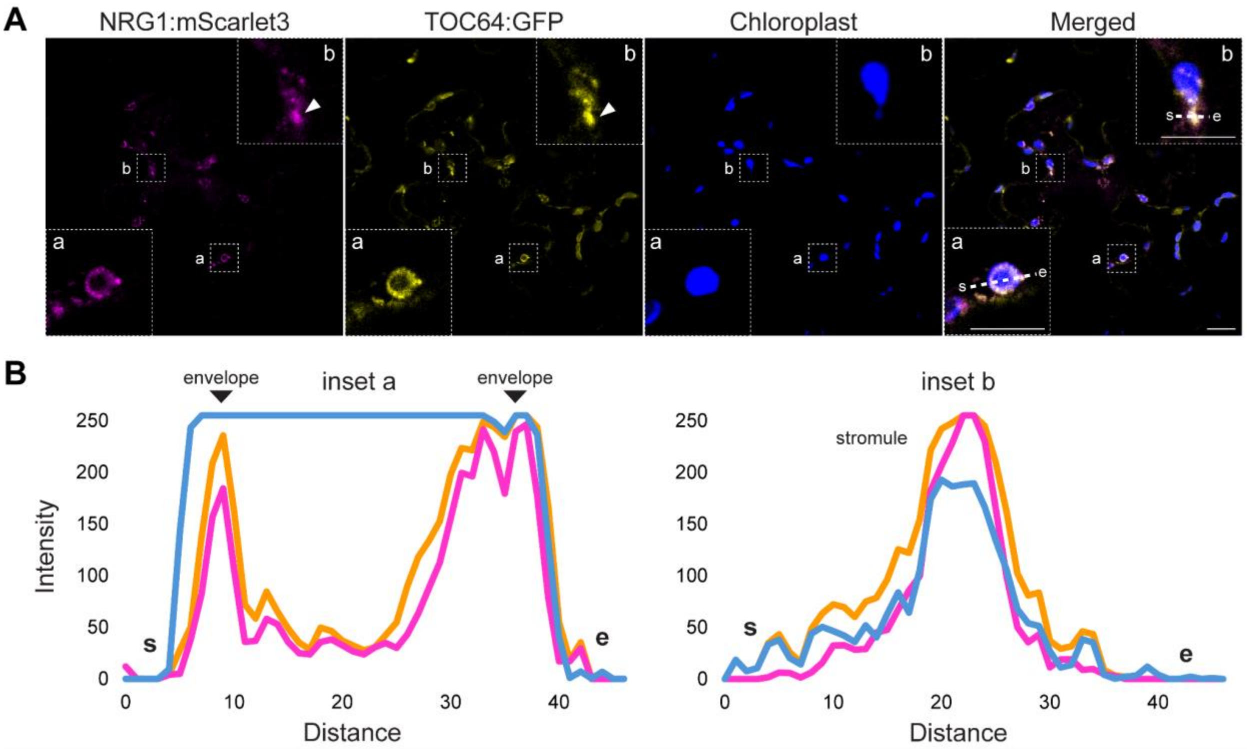


**Fig. S6 NbNRG1 co-localizes with chloroplast outer-membrane protein TOC64.** **(A)** Confocal micrograph of *N. benthamiana* leaf epidermal cells transiently expressing NRG1:mScarlet3 and TOC64:GFP. Chloroplasts are visualized using auto-fluorescence at 648-709 nm. Dashed lines across panels correspond to insets a and b. Inset b highlights NbNRG1 targeting of stromules. All images shown are single-plane images. Scale bars represent 10 µm. Lines (s to e) in the inset panels – a and b - correspond to **(B)** line intensity plots depicting the relative fluorescence across the marked area. Raw fluorescence intensity data can be found in Data S2. Imaging was done with at least two separate leaf patches per panel.

**Fig. S7**

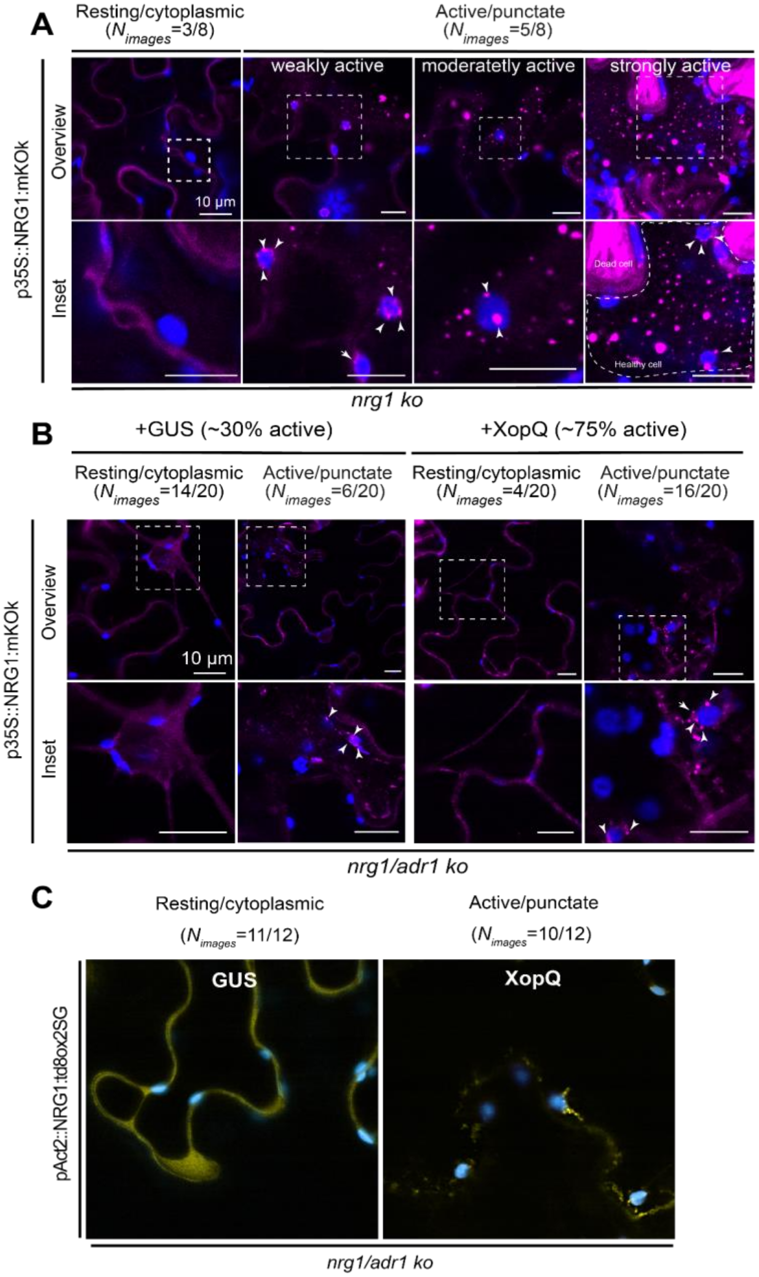


**Fig. S7 Alternative fluorescent constructs produced weaker cell death allowing resting state imaging of full-length NbNRG1.**
(**A**) Confocal micrographs of *N. benthamiana nrg1* KO leaf epidermal cells transiently expressing 35S::NRG1:mKOK. Images shown are single-plane images. Dashed lines in overview correspond to inset. Arrows indicate examples of puncta around chloroplasts. Scale bars represent 10 µm. Imaging was done with a minimum of two leaf patches per condition. For weakly, moderate and strongly active panels, the number of puncta is used as a reference. The highlighted area in the inset panel for the strongly active state reflects a healthy cell of interest. Dead cells are labelled clearly. (**B**) Confocal micrographs of *N. benthamiana nrg1/adr1* double KO leaf epidermal cells transiently expressing 35S::NRG1:mKOK with XopQ or GUS control. Images shown are single-plane images. Dashed lines in overview correspond to inset. Arrows indicate examples of puncta around chloroplasts. Scale bars represent 10 µm. For panels A and B, NRG1:mKOK is colored as

a magenta signal and the chloroplast auto-fluorescene is shown as blue. (**C**) Confocal micrographs of *N. benthamiana* leaf epidermal cells transiently expressing pAct2::NRG1:td8ox2SG with XopQ or GUS control. NRG1:td8ox2SG is shown as yellow and chloroplast auto-fluorescence as blue. Images shown are single-plane images. Scale bar represents 10 µm. Imaging was done with at least two separate leaf patches per panel.

**Fig. S8.**

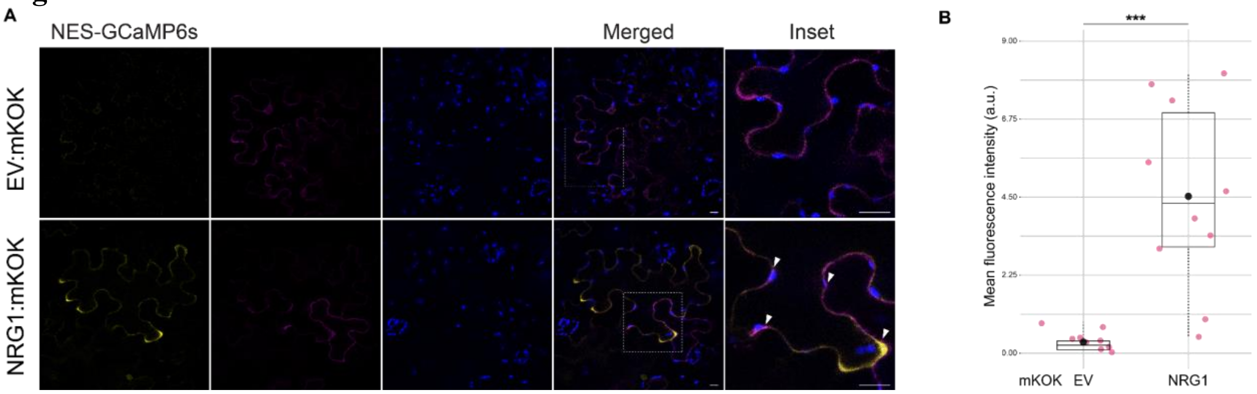


**Fig. S8 NbNRG1 expression leads to calcium accumulation in the cytoplasm.**
(**A**) Confocal micrographs of *N. benthamiana* wild type leaf epidermal cells transiently expressing NbNRG1:mKOK or mKOK control and an GFP-labelled cytoplasmic $Ca^{2+}$ sensor (NES-GCaMP6s:GFP). Images shown are single-plane images. Chloroplasts are visualized using auto-fluorescence at 648-709 nm. Dashed lines in merged panels correspond to insets. Scale bars represent 10 µm. Imaging was done with three separate leaf patches at 28-hours post agroinfiltration. White arrows indicate NbNRG1:mKOK puncta. (**B**) Box and dot plot showing the mean fluorescence intensity for NES-GCaMP6s:GFP when co-expressed with either EV:mKOK or NbNRG1:mKOK. Each cell is highlighted with a pink dot. The mean value for each condition is shown as a black dot. Asterisks indicate statistically significant difference of NRG1:mKOK co-expression with NES-GCaMP6s:GFP compared to the control, EV:mKOK (NRG1:mKOK – Wilcoxon, W=2, p-value<0.005). Raw fluorescence intensity data can be found at Data S2.

**Fig. S9**

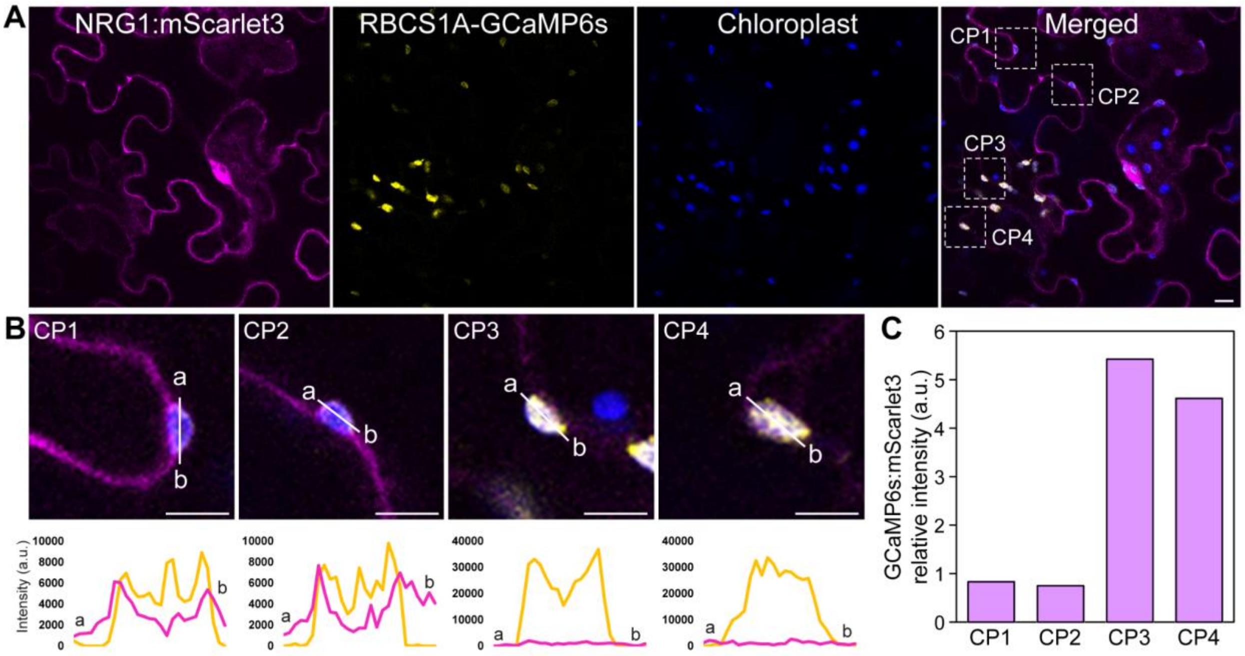


**Fig. S9 NRG1 depletes calcium stores of chloroplasts.**
(**A**) Confocal micrograph of *N. benthamiana* leaf epidermal cells transiently expressing NRG1:mScarlet3 and RBCS1A-GCaMP6s:GFP. Chloroplasts are visualized using auto-fluorescence at 648-709 nm. Dashed lines in the merged panel correspond to insets shown in (B). Imaging was done with at least two separate leaf patches per panel. (**B**) Insets from the merged panel in (A) focusing on chloroplasts (CP). Lines (a to b) in the inset panels - CP1 to 4 - correspond to line intensity plots depicting the relative fluorescence across the marked area. (**C**) Relative intensity of GCaMP6s:mScarlet3 for each chloroplast (CP1-4) highlighted in (B).

**Fig. S10**

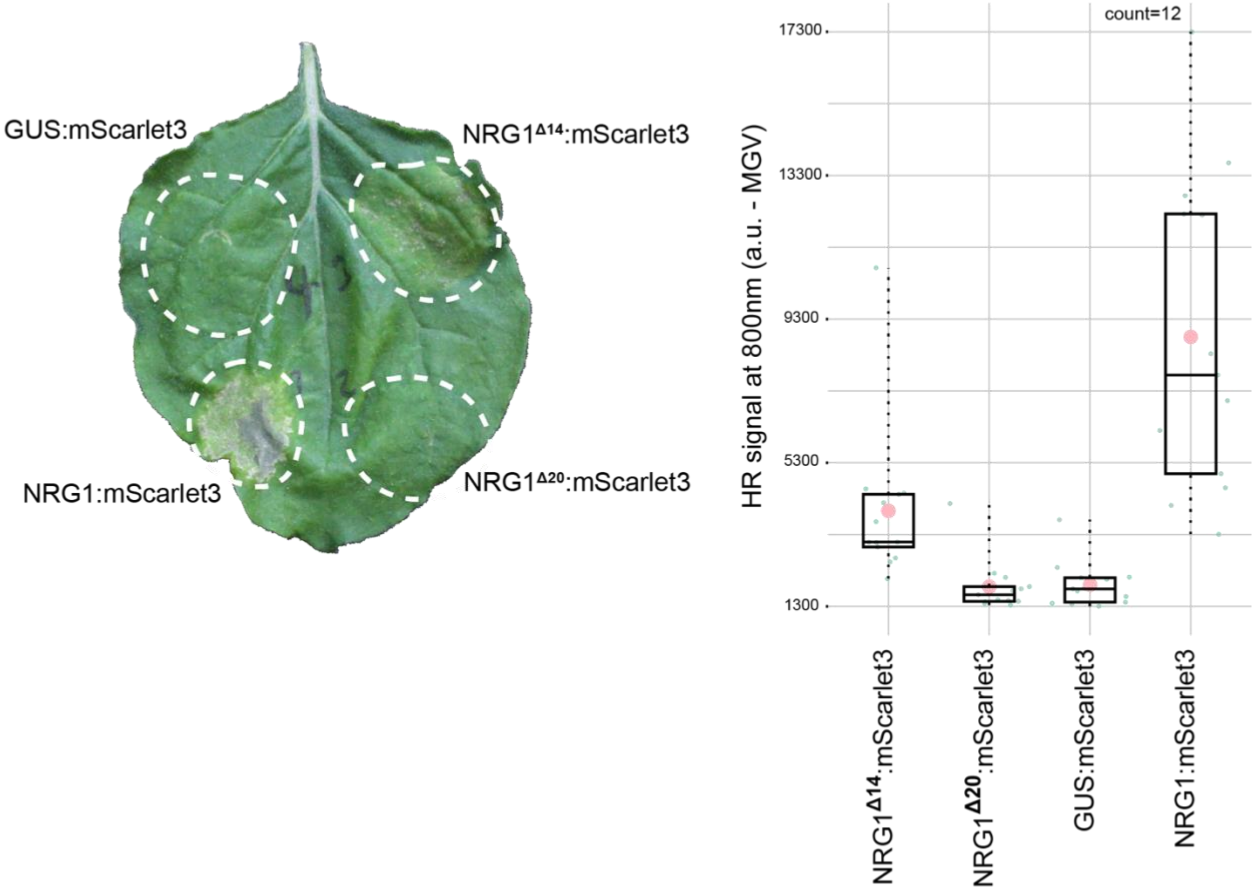


**Fig. S10 20 amino acid truncation of NbNRG1 hampers cell death.**
A representative leaf picture from *N. benthamiana* wild-type plants showing HR after expression of NRG1$^{\Delta14}$:mScarlet3, NRG1:mScarlet3, NRG1$^{\Delta20}$:mScarlet3 or GUS:mScarlet3. Box and dot plot showing the relative HR signal at 800 nm (infrared) for 12 leaves. Each leaf is highlighted with a green dot and the mean value for that condition in a larger pink dot. Raw HR intensity can be found at Data S3.

**Fig. S11**

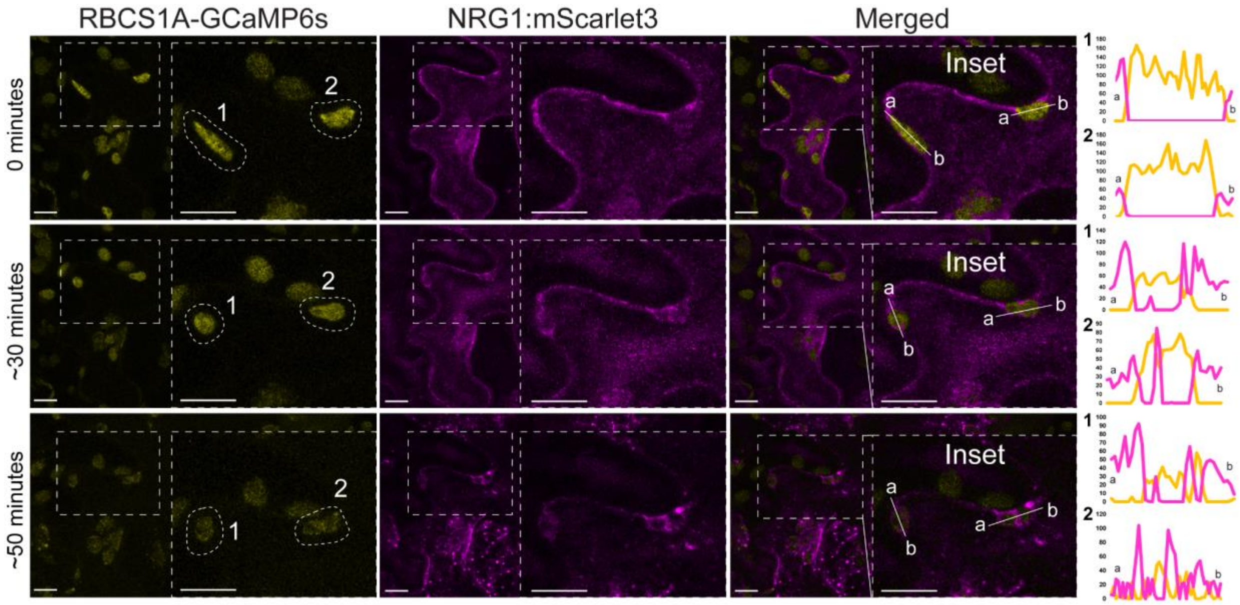


**Fig. S11 Formation of NRG1 puncta on the chloroplast envelope is followed by calcium depletion from the chloroplast.**

Confocal micrographs of *N. benthamiana* wild-type leaf epidermal cells transiently expressing NRG1:mScarlet3 and RBCS1A-GCaMP6s:GFP. Images shown are single-plane images from a time series collected over an hour time-period. Dashed lines around chloroplasts – 1 and 2 – correspond to tracked chloroplasts over time and inset. Lines (a to b) in the insets correspond to line intensity plots depicting the relative fluorescence across the marked area.

**Fig. S12**

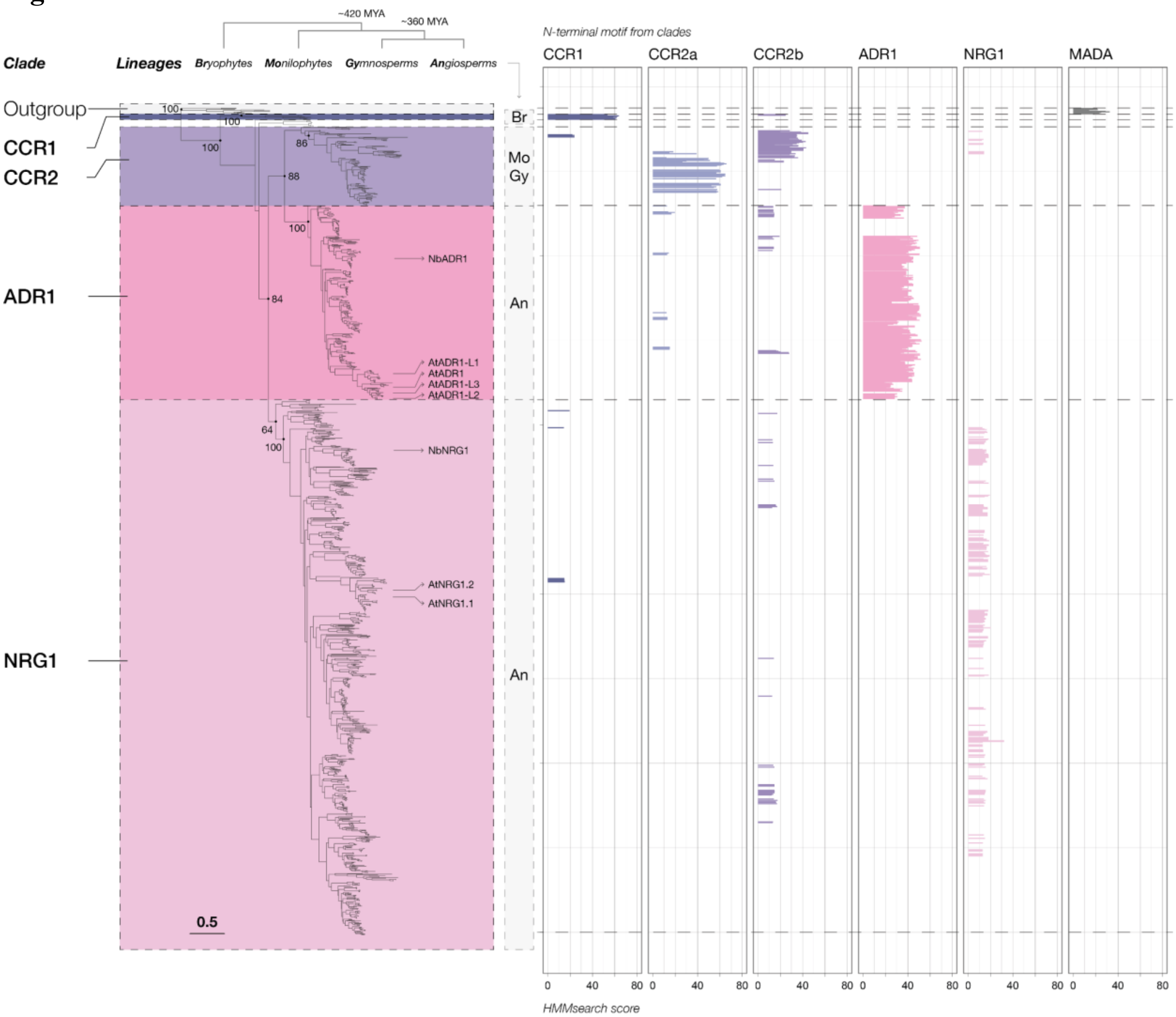


**Fig. S12 CC$_R$-NLRs exhibit distinct phylogenetic characteristics in their coiled-coil domains.** The phylogenetic tree illustrates the relationships among 56,280 CC$_R$-NLRs based on their NB-ARC domain sequences. Four well-supported clades are highlighted: the CCR1 and CCR2 clades (purple), the ADR1 clade (pink) and the NRG1 clade (lilac). Distinct N-terminal motifs in the CC$_R$ domains of each clade were identified using MEME analysis and their HMMsearch scores were plotted with the CC-NLR MADA motif as a control (right). The tree is rooted with sequences from AtZAR1, NbZAR1, and NRCs from tomato as the outgroup.

**Fig. S13**

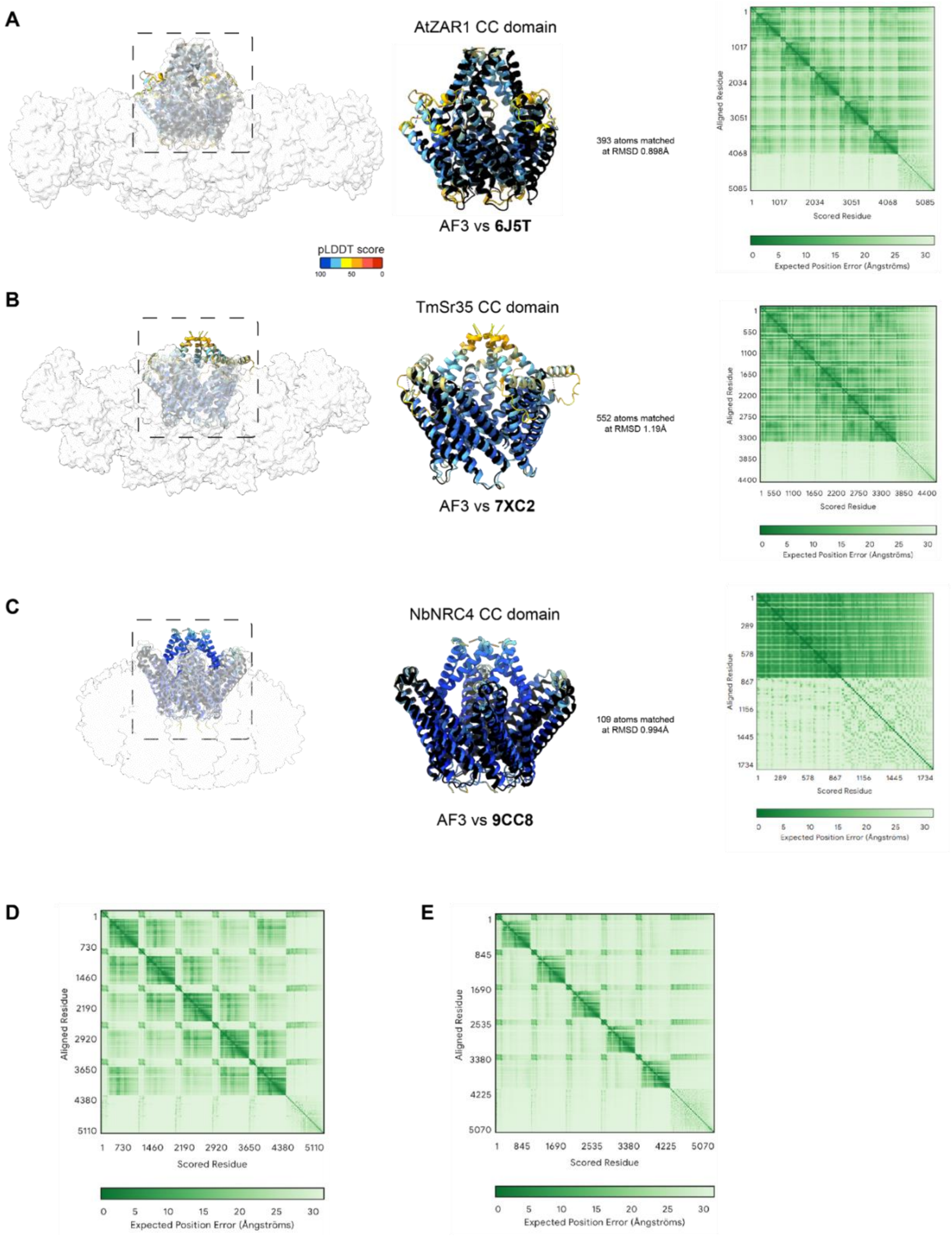
A
AtZAR1 CC domain
393 atoms matched
at RMSD 0.898Å
AF3 vs 6J5T
pLDDT score
100
50
0
Aligned Residue
Scored Residue
Expected Position Error (Ångströms)
B
TmSr35 CC domain
552 atoms matched
at RMSD 1.19Å
AF3 vs 7XC2
Aligned Residue
Scored Residue
Expected Position Error (Ångströms)
C
NbNRC4 CC domain
109 atoms matched
at RMSD 0.994Å
AF3 vs 9CC8
Aligned Residue
Scored Residue
Expected Position Error (Ångströms)
D
Aligned Residue
Scored Residue
Expected Position Error (Ångströms)
E
Aligned Residue
Scored Residue
Expected Position Error (Ångströms)

**Fig. S13 Quality analysis of AlphaFold3 predicted resistosome and coiled-coil domain structures.**
(**A**) Coiled-coil domain structure (black) and AlphaFold3 prediction (colored by pLDDT) overlaid and fitted into the cryoEM density map of AtZAR1 (6J5T) with an RMSD of 0.898Å. The PAE plot for AlphaFold3 prediction of AtZAR1 resistosome was downloaded from the AlphaFold3 server. (**B**) Coiled-coil domain structure (black) and AlphaFold3 prediction (colored by pLDDT) overlaid and fitted into the cryoEM density map of TmSr35 (7XC2) with an RMSD of 1.19Å. The PAE plot for AlphaFold3 prediction of TmSr35 resistosome was downloaded from the AlphaFold3 server. (**C**) Coiled-coil domain structure (black) and CC domain AlphaFold3 prediction (colored by pLDDT) overlaid and fitted into the cryoEM density map of NbNRC4 (9CC8) with an RMSD of 0.994Å. The PAE plot for AlphaFold3 prediction of NbNRC4 CC domain was downloaded from the AlphaFold3 server. (**D**) The PAE plot for AlphaFold3 prediction of NbNRG1 resistosome was downloaded from the AlphaFold3 server. (**E**) The PAE plot for AlphaFold3 prediction of NbADR1 resistosome was downloaded from the AlphaFold3 server. AF3 models can be found at Data S13.

**Fig. S14**

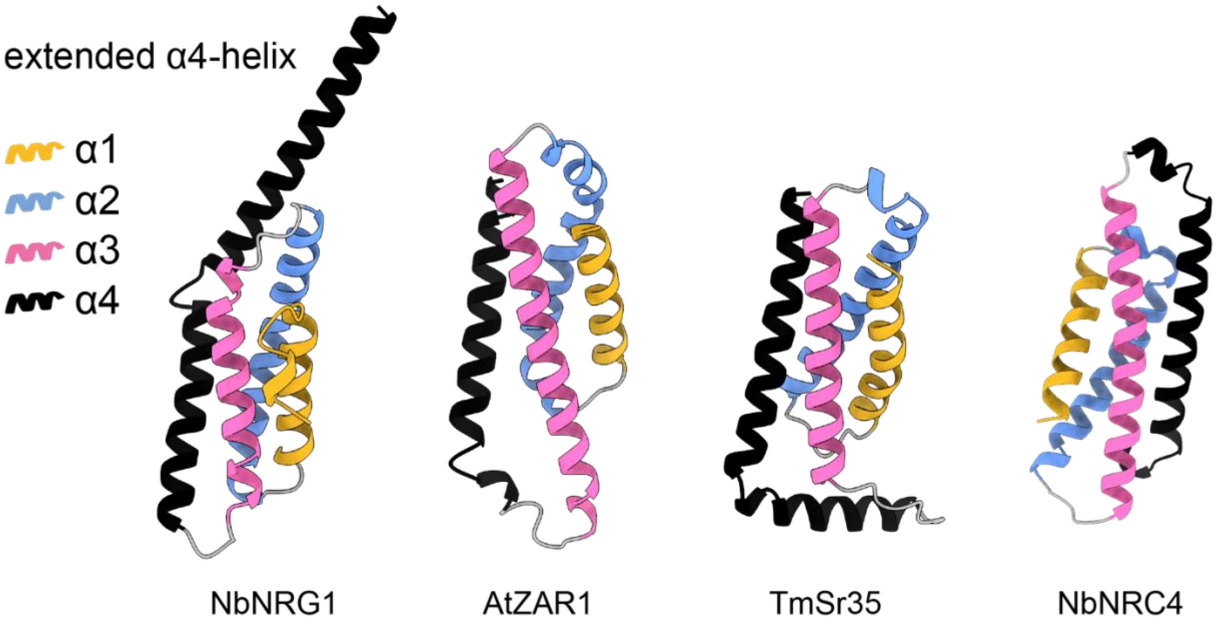


**Fig. S14 CC$_R$-NLR NbNRG1 has an extended fourth alpha-helix in the coiled-coil domain compared to CC-NLRs NbNRC4, AtZAR1 and TmSr35.**
Alpha-helix one is colored yellow, two blue, three pink and four black in all presented models.

**Fig. S15**

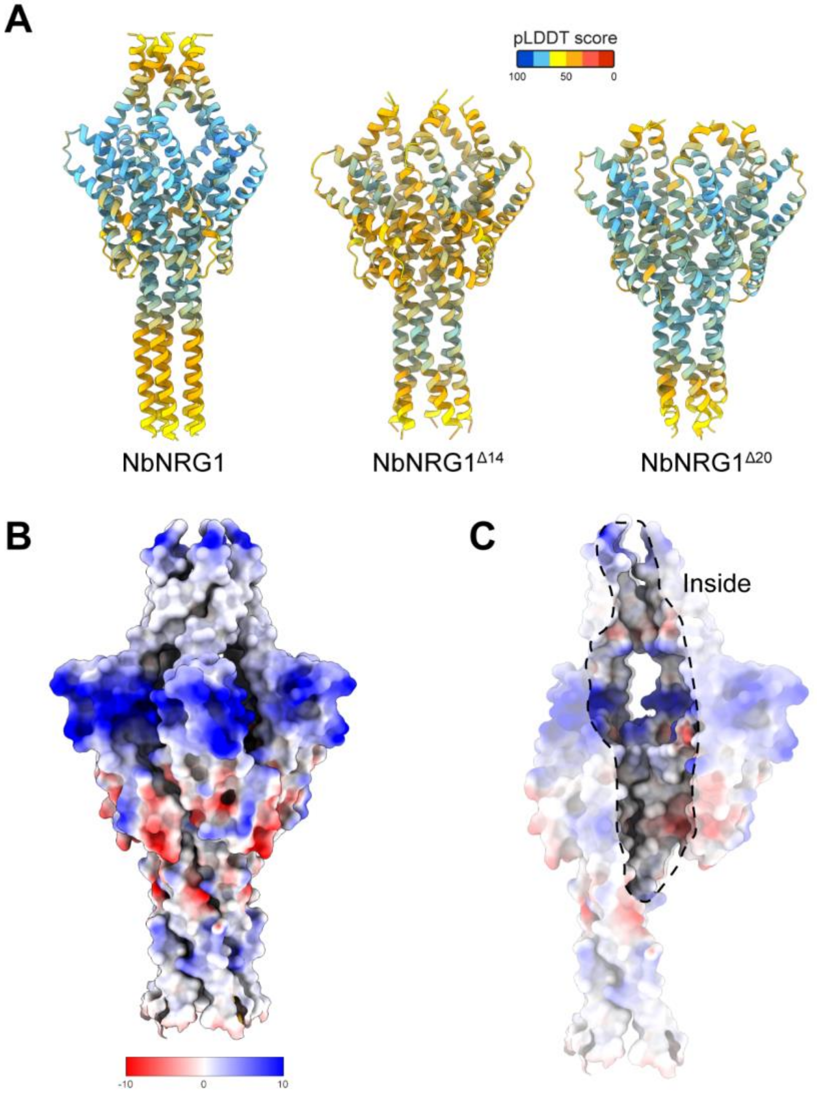


**Fig. S15 AlphaFold3 modelling of NRG1 and truncates.**
(**A**) NbNRG1 was modelled using AlphaFold 3 alongside its truncates missing the first 14 and 20 amino acids to illustrate their ability to form shorter oligomeric structures. Predicted structures are colored based on their pLDDT scores. (**B**) and (**C**) Electrostatic potential of the predicted NbNRG1 CC domain structure. The model is colored based on electrostatic potential, where blue indicates hydrophilic and red indicates hydrophobic regions. The inside of the CC pore structure is highlighted in (C).

**Fig. S16**

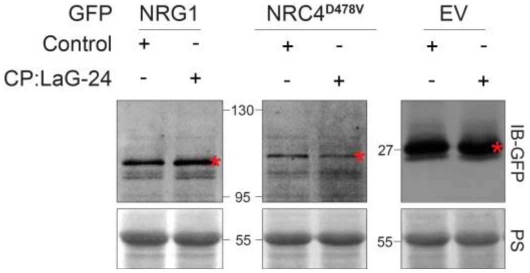


**Fig. S16 Nanobody-enrichment at the chloroplast does not significantly alter soluble protein levels.**

Protein levels for NRG1:GFP, NRC4$^{D478V}$:GFP or EV:GFP control with and without CP:LaG-24 nanobody. Total protein extracted at 26-hours post infiltration from three different leaves.

**Fig. S17**

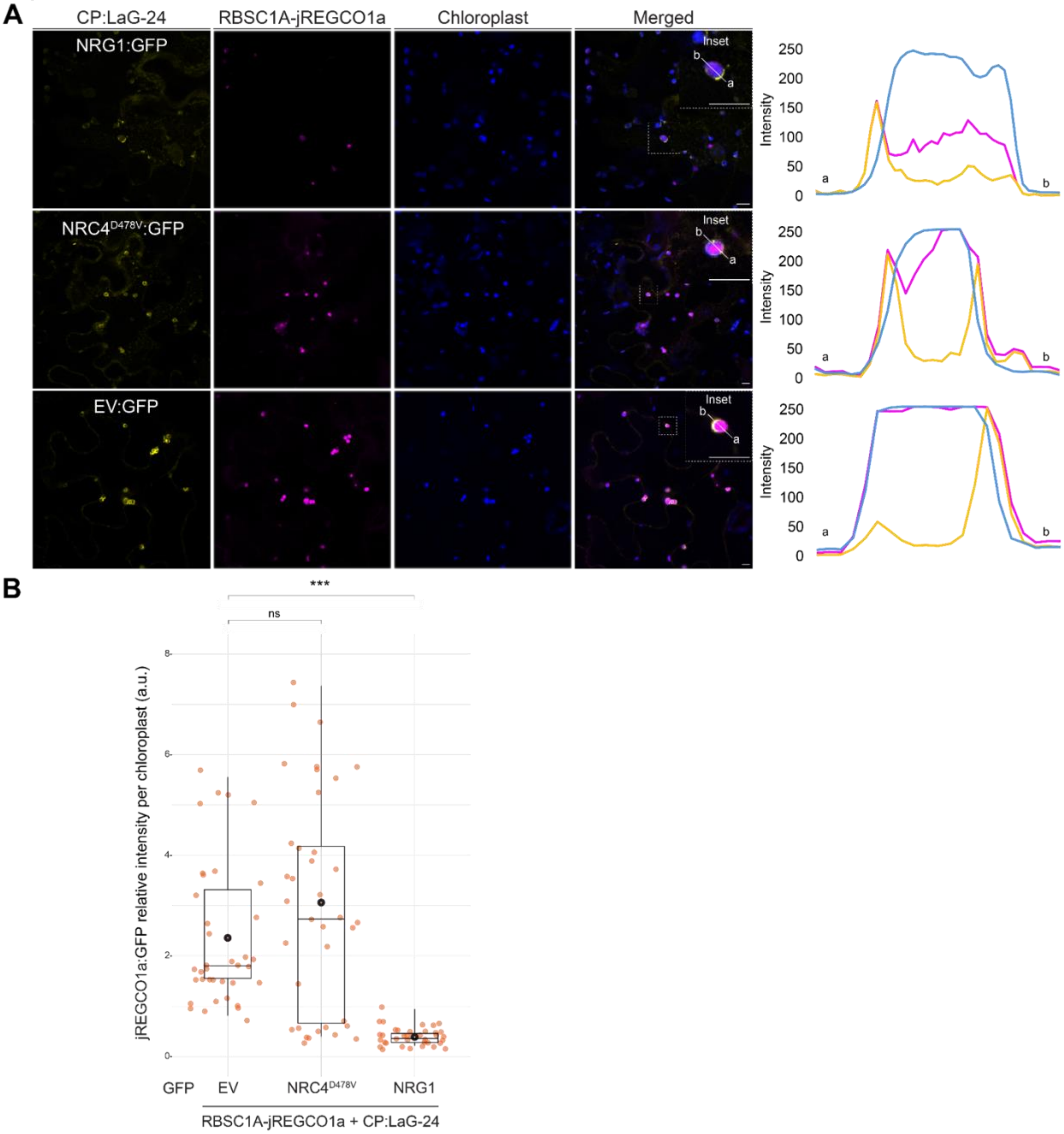


**Fig. S17 NbNRC4 cannot facilitate calcium ion channeling from the chloroplast membrane.** (**A**) Confocal micrographs of *N. benthamiana* wild type leaf epidermal cells transiently expressing NbNRC4$^{D478V}$:GFP, NbNRG1:GFP or GFP control with the nanobody CP:LaG-24 and an RFP-labelled $Ca^{2+}$ sensor (RBSC1A-jREGCO1a). Images shown are single-plane images. Dashed lines in the merged panel correspond to inset. Lines (a to b) in the insets correspond to line intensity plots depicting the relative fluorescence across the marked area. Chloroplasts are visualized using auto-fluorescence at 648-709 nm. Scale bars represent 10 µm. Imaging was done with at least two separate leaf patches per panel. (**B**) Box and dot plot showing the jREGCO1a(RFP):GFP relative intensity per chloroplast for 37 chloroplasts from two separate leaf patches per sample. Each chloroplast is highlighted with an orange dot. The mean value for each condition is shown as a black dot. Asterisks indicate statistically significant difference of NRG1:GFP or NRC4$^{D478V}$:GFP co-expression with RBSC1A-jREGCO1a and CP:LaG-24 compared to the control, EV:GFP (NRC4$^{D478V}$:GFP – Wilcoxon, W=585, p-value=0.2844 and NRG1:GFP – Wilcoxon, W=1367, p-value<0.005). Raw fluorescence intensity data can be found at Data S2.

**Fig. S18**

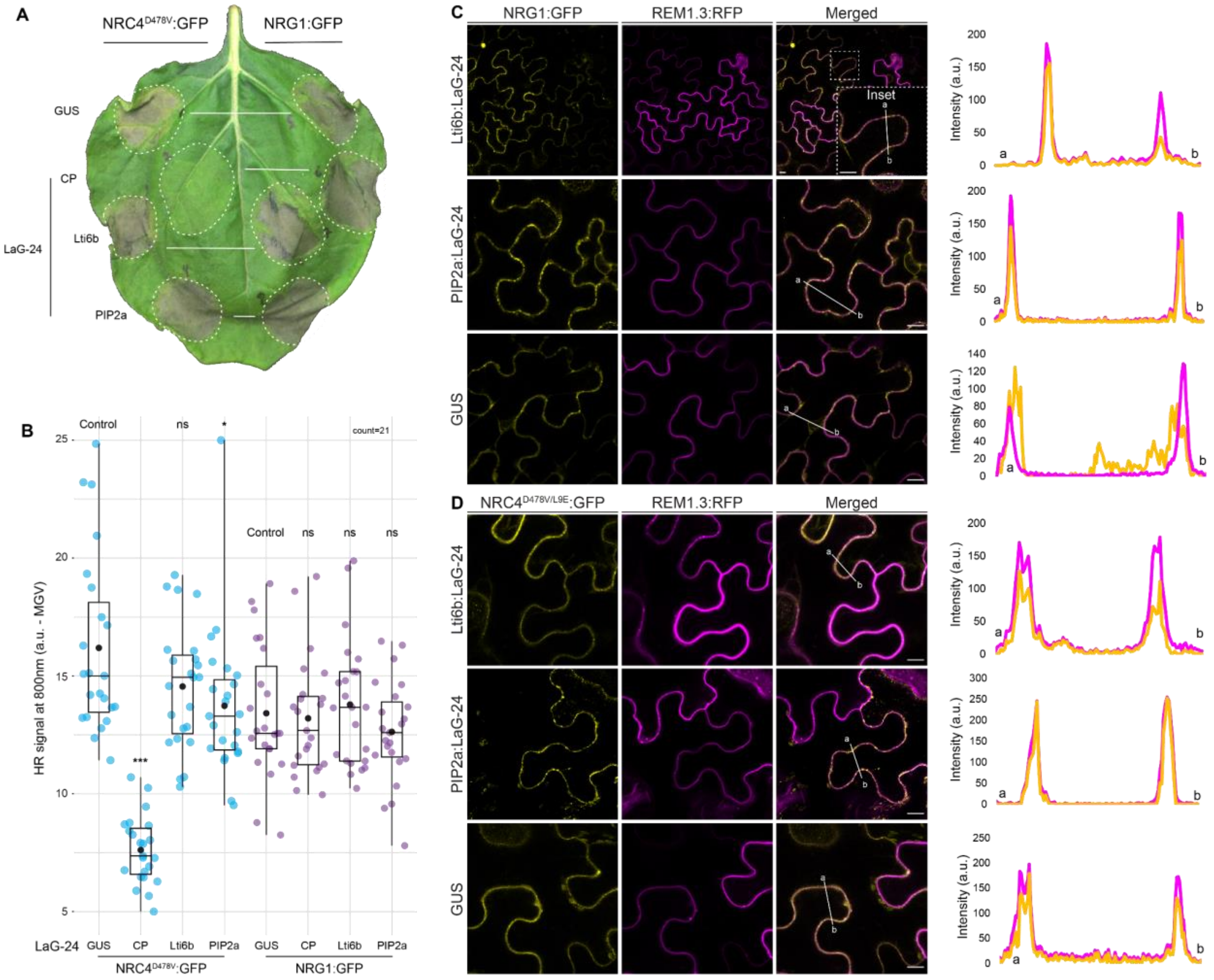


**Fig. S18 NbNRG1 and NbNRC4 retain HR activity under partial plasma membrane trapping.**

(**A**) A representative leaf picture from *N. benthamiana* wild-type plants showing HR after expression of NRC4$^{D478V}$:GFP and NRG1:GFP with CP:LaG-24, Lti6b:LaG-24, PIP2a:LaG-24 or GUS control. (**B**) Box and dot plot showing the relative HR signal at 800 nm (infrared) for 21 leaves. Each leaf is highlighted with a blue dot for NRC4$^{D478V}$ and purple dot for NRG1. The mean value for each condition is shown in a larger back dot. Asterisks indicate statistically significant difference of NRG1:GFP and NRC4$^{D478V}$:GFP co-expression with different nanobodies compared to the control, GUS (NRC4$^{D478V}$:GFP + CP:LaG-24 – Wilcoxon, W=529, p-value= 2.429e-13. NRC4$^{D478V}$:GFP + Lti6b:LaG-24 – Wilcoxon, W=322, p-value= 0.2127. NRC4$^{D478V}$:GFP + PIP2a:LaG-24 – Wilcoxon, W=374, p-value= 0.01561. NRG1:GFP + CP:LaG-24 – t-test, t=0.274, p-value= 0.7855. NRG1:GFP + Lti6b:LaG-24 – t-test, t=-0.453, p-value=0.6528. NRG1:GFP + PIP2a:LaG-24 – t-test, t=1.06, p-value= 0.2951). Raw HR intensity can be found at Data S3. (**C**) and (**D**) Confocal micrographs of *N. benthamiana* wild type leaf epidermal cells transiently expressing (C) NbNRG1 and (D) NbNRC4$^{D478V/L9E}$:GFP with the nanobodies Lti6b:LaG-24, PIP2a:LaG-24 or GUS control and REM1.3:RFP as a plasma membrane marker. Images shown are single-plane images. Dashed lines in merged panel correspond to inset. Lines (a to b) correspond to line intensity plots depicting the relative fluorescence across the marked area. Scale bars represent 10 µm.

**Fig. S19**

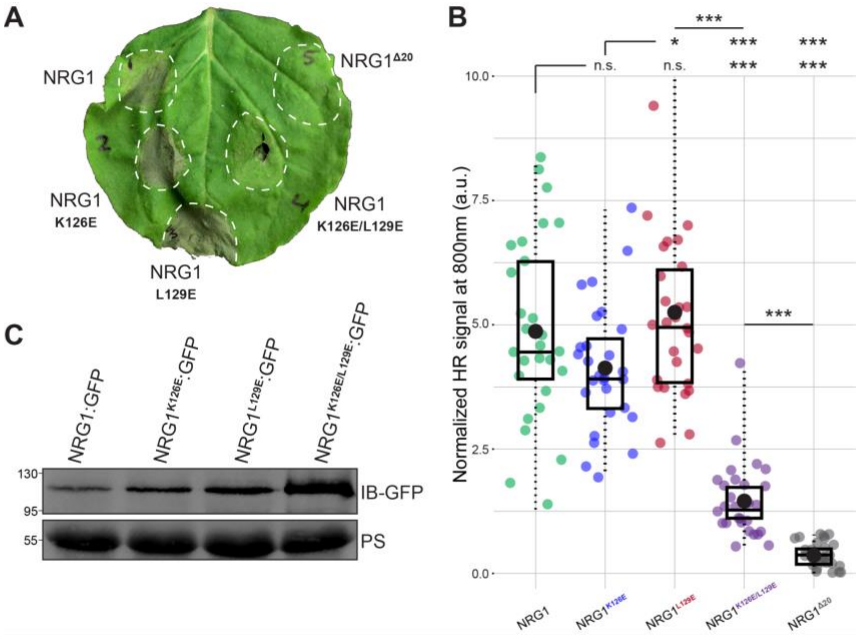


**Fig. S19 Alpha-helix four residues K126 and L129 are essential for NbNRG1 HR execution.** **(A)** A representative leaf picture from wild-type *N. benthamiana* plants showing HR after expression of NbNRG1:GFP, NbNRG1$^{K126E}$:GFP, NbNRG1$^{L129E}$:GFP, NbNRG1$^{K126E/L129E}$:GFP or NRG1$^{\Delta20}$:GFP (negative control – no HR). (**B**) Box and dot plot showing the normalized HR signal at 800 nm (infrared) for 26 leaves from two different replicates. Each leaf is highlighted with a green dot for NRG1:GFP, blue dot for NbNRG1$^{K126E}$:GFP, red dot for NbNRG1$^{L129E}$:GFP, purple dot for NbNRG1$^{K126E/L129E}$:GFP and a gray dot for NRG1$^{\Delta20}$:GFP. The mean value for each condition is shown as a black dot. Three asterisks indicate a statistically significant difference of p-value<0.005. Raw HR intensity can be found at Data S3. **(C)** Protein levels for NbNRG1:GFP, NbNRG1$^{K126E}$:GFP, NbNRG1$^{L129E}$:GFP and NbNRG1$^{K126E/L129E}$:GFP. Total protein extracted at 26 to 28-hours post agroinfiltration.

**Fig. S20**

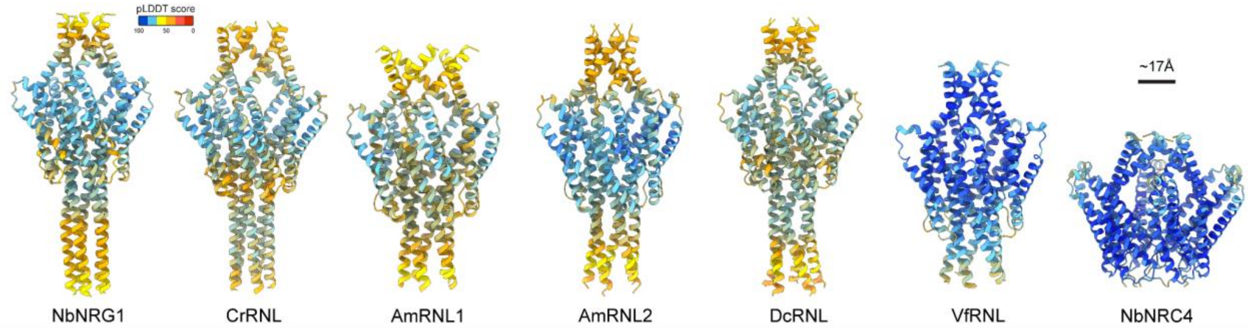


**Fig. S20 AlphaFold 3 predicted models for NRG1-like $CC_R$-NLRs studied.**

NbNRG1 and variants studied in Fig. 4 were modelled using AlphaFold 3 and colored based on their respective pLDDT metric. Scale bar represents 17 angstroms.

**Fig. S21**

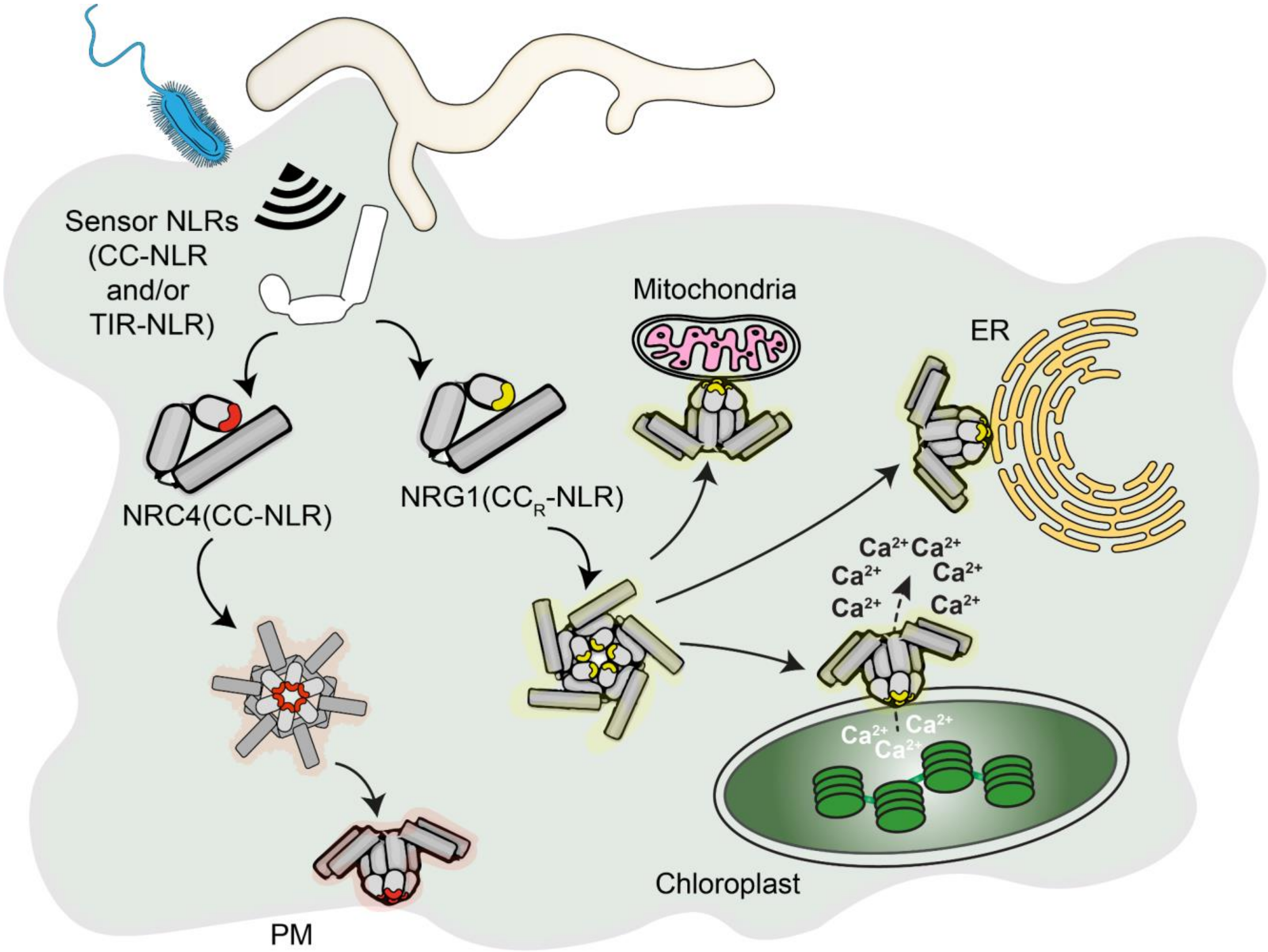


**Fig. S21 A representative model for differential NLR localization.**

Upon pathogen recognition, NRC4 targets the plasma membrane, whereas NRG1 can target various intracellular membrane compartments and channel calcium ions from the chloroplast. Endoplasmic reticulum; ER, plasma membrane; PM.

**Table S1. List of all constructs used in this research**

| Construct | Plasmids | Reference |
|---|---|---|
| p35S::NbNRG1:GFP (with p19) | pJK268c_mRFP1, pICH51288, pJK-0-320, pICSL50034, pICH41414 | This work |
| p35S::NbNRG1:GFP (without p19) | pJK001c_mRFP1, pICH51288, pJK-0-320, pICSL50034, pICH41414 | This work |
| p35S::NbNRG1:mKOK (without p19) | pJK001c_mRFP1, pICH51288, pJK-0-320, pICH41414 | This work |
| p35S::NbNRG1$^{\Delta14}$:GFP (with p19) | pJK268c_mRFP1, pICH51288, pJK-0-321, pICSL50034, pICH41414 | This work |
| p35S::NbNRG1$^{\Delta14}$:GFP (without p19) | pJK001c_mRFP1, pICH51288, pJK-0-321, pICSL50034, pICH41414 | This work |
| p35S::EV:GFP (with p19) | pJK268c_mRFP1, pICH51288, pICSL50034, pICH41414 | This work |
| p35S::EV:GFP (without p19) | pJK001c_mRFP1, pICH51288, pICSL50034, pICH41414 | This work |
| 2xp35S::XopQ | pJK001c_mRFP1, pICH51288, pUC57-Kan-XopQ, pICH41414 | This work |
| NRC4$^{L9E/DV}$:GFP | / | *(26)* |
| FLAG:GUS | / | *(75)* |
| RFP:EV | / | *(59)* |
| RFP:ER (SP:RFP:HDEL) | / | *(76, 77)* |

| | | |
|---|---|---|
| Mitochondria:mCherry ($ScCOX4_{1-29}$:mCherry) | / | *(75, 77, 78)* |
| CTP1:mCherry | / | *(33)* |
| Golgi:mCherry ($GmMan1_{1-49}$:mCherry) | / | *(75, 77, 79)* |
| RFP:Remorin1.3 | / | *(80)* |
| TOC64::GFP | / | *(34)* |
| p*At*Act2::NbNRG1$^{\Delta14}$:GFP (without P19) | pJK001c_mRFP1, pICH87644, pJK-0-321, pICSL50034, pICH41414 | This work |
| p35S::NbNRG1:td8ox2StayGold (without p19) | pJK001c_mRFP1, pICH51288, pJK-0-320, td8ox2StayGold-pTwist_Amp_HC (synthesized, Twist Bioscience), pICH41414 | This work |
| p35S::NbNRG1:mScarlet3 (without p19) | pJK001c_mRFP1, pICH51288, pJK-0-320, pJK-CT-0005, pICH41414 | This work |
| p35S::NbNRG1$^{\Delta20}$:mScarlet3 (without p19) | pJK001c_mRFP1, pICH51288, pFK1S-0240, pJK-CT-0005, pICH41414 | This work |
| p35S::GUS:mScarlet3 (without p19) | pJK001c_mRFP1, pICH51288, GUS-pTwist_Amp_HC (synthetized, Twist Bioscience), pJK-CT-0005, pICH41414 | This work |
| p35S::RBCS1A-GCaMP6s | pJK001c_mRFP1, pICH51288, pJK-NT-0001, pJK-CDS-0110, pICSL50028, pICH41414 | This work |
| p35S::NES-GCaMP6s | pJK001c_mRFP1, | This work |

| | | |
|---|---|---|
| | pICH51288, pJK-NT-0018, pJK-CDS-0110, pICSL50028, pICH41414 | |
| p35S 5U::NbNRG1:mTurquoise2::tHSP (no p19) | pJK001c_mRFP1, pICH51266, pJK-0-320, pJK-CT-0007, pICH41414 | This work |
| p35S 5U::NbNRG1:mGold:tHSP (no p19) | pJK001c_mRFP1, pICH51266, pJK-0-320, pJK-CT-0009, pICH41414 | This work |
| p35S::RCaMP1h | / | *(40)* |
| CP-LaG-24 (p35S::NtHR-LaG24) | pJK001c_mRFP1, pICSL13008, pJK-CDS-0007, pJK-NT-0027, pICSL50028, pICSL60008 | This work |
| p35S::Lti6b-LaG-24 | pJK001c_mRFP1, pICSL13008, pJK-CDS-0007, pJK-NT-0030, pICSL50028, pICSL60008 | This work |
| p35S::PIP2a:LaG-24 | pJK001c_mRFP1, pICSL13008, pJK-CDS-0007, pJK-NT-0031, pICSL50028, pICSL60008 | This work |
| p35S 5U::NbNRG1:GFP::tHSP (no p19) | pJK001c_mRFP1, pICH51266, pJK-0-320, pICSL50034, pICSL60008 | This work |
| p35S 5U::NbNRG1 $^{K126E}$:GFP::tHSP (no p19) | pJK001c_mRFP1, pICH51266, pFK1S-0243, pICSL50034, pICSL60008 | This work |
| p35S 5U::NbNRG1 $^{L129E}$:GFP::tHSP (no p19) | pJK001c_mRFP1, pICH51266, pFK1S-0244, pICSL50034, pICSL60008 | This work |
| p35S 5U::NbNRG1 $^{K126E/L129E}$:GFP::tHSP (no p19) | pJK001c_mRFP1, pICH51266, pFK1S-0330, pICSL50034, pICSL60008 | This work |

| | | |
|---|---|---|
| CrCC$_{R}$-NLR$^{D559V}$:YFP | pICH47742, pICH85281, Ceric.39G035100.1_DV (synthesized, Genewiz), pICSL50005, pICSL60008 | This work |
| AmCC$_{R}$-NLR1$^{D478V}$:YFP | pICH47742, pICH85281, AmTrH2.13G128800.1_DV (synthesized, Genewiz), pICSL50005, pICSL60008 | This work |
| AmCC$_{R}$-NLR2$^{D475V}$:YFP | pICH47742, pICH85281, AmTrH2.13G128900.1_DV (synthesized, Genewiz), pICSL50005, pICSL60008 | This work |
| p35S::DcCC$_{R}$-NLR:GFP | DcRNL PCR product, pK7WGF2 | This work |
| p35S::VfCC$_{R}$-NLR:GFP | pJK268c_mRFP1, pICH51288, pFK1S-0174, pICSL50034, pICH41414 | This work |

**Table S2. List of oligos used for cloning in this study**

| | |
|---|---|
| **Oligos for generating the NbNRG1$^{\Delta 20}$ mutant** | |
| **Name** | **Sequence** |
| NbNRG1 d20 F1 | GCTGTGCTGGACGTGGG |
| NbNRG1 d20 R2 | CATTTGAGACCACAGAGTGATTAATGAATC |
| **Oligos for generating NbNRG1$^{K126E}$ mutant** | |
| **Name** | **Sequence** |
| NbNRG1 K126E F | CAGGGACTCTGAAATCATCCTGGT |
| NbNRG1 K126E R | CACACCTGAATAAAGCCGTG |
| **Oligos for generating NbNRG1$^{L129E}$ mutant** | |
| **Name** | **Sequence** |
| NbNRG1 L129E F | TAAGATCATCGAAGTGAATGTGATCGAGCACGG |
| NbNRG1 L129E R | GAGTCCCTGCACACC |
| **Oligos for generating NbNRG1$^{K126E/L129E}$ mutant** | |
| **Name** | **Sequence** |
| NbNRG1 K126E, L129E F | TTGAAGACAAATCGAGGTGAATGTGATCGAGCACGGCAAG |
| NbNRG1 K126E, L129E R | TTGAAGACAACGATGATCTCAGAGTCCCTGCACACCTG |
| **Oligos for cloning DcRNL into pK7WGF2** | |
| **Name** | **Sequence** |
| DcNRG1 GA F1 | CAGGCGGCCGCACTAGTGATATGGATCTTATTGGTGGTGG AATTATTG |
| DcNRG1 GA R2 | GCAGATCCAGCAGATCCGATATGATTAAGCCAATTAAGAT TAATATCTTCCTTAAC |
| **Oligos for cloning VfRNL into pICSL01005CP** | |
| **Name** | **Sequence** |
| VfNRG1-like GG F5 | CACTCTGTGGTCTCAAATGGCAGGGTTGGTTGAAGG |
| VfNRG1-like GG R5 | ATTCGTGGTCTCACGAACCTACACTAATGAATAAGTTAATT TCAGCCTCC |
